# Mitochondrial reactive oxygen species cause arrhythmias in hypertrophic cardiomyopathy

**DOI:** 10.1101/2024.10.02.616214

**Authors:** Michael Kohlhaas, Vasco Sequeira, Shan Parikh, Alexander Dietl, Olga Richter, Johannes Bay, Edoardo Bertero, Julia Schwemmlein, Qinghai Tian, Felix W. Friedrich, Frederik Flenner, Alexander G. Nickel, Felix Alscher, Maithily S. Nanadikar, Raghav Venkataraman, Franz J. Baudenbacher, Reinhard Kappl, Victoria Johnson, Leticia Prates Roma, Andrej Kasakow, Mathias Hohl, Ulrich Laufs, Markus Hoth, Dörthe M. Katschinski, Michael Böhm, Peter Lipp, Lucie Carrier, Thomas Eschenhagen, Björn C. Knollmann, Christoph Maack

**Affiliations:** Comprehensive Heart Failure Center Würzburg, University Clinic Würzburg, Germany; (former) affiliation, where part of the experiments were performed; Clinic for Internal Medicine III, University Clinic of the Saarland, Homburg, Germany; Center for Arrhythmia Research and Therapeutics, Department of Medicine, Vanderbilt University Medical Center, Nashville, TN, USA; Department of Internal Medicine II, University Hospital Regensburg, Regensburg, Germany; Institute for Cell Biology, Saarland University, Homburg, Germany; Department of Experimental Pharmacology and Toxicology, Cardiovascular Research Center, University Medical Center Hamburg-Eppendorf, Hamburg, Germany; DZHK (German Centre for Cardiovascular Research), partner site Hamburg/Kiel/Lübeck, Germany; Department of Cardiovascular Physiology, University Medical Center Göttingen, Göttingen; Department of Biomedical Engineering, Vanderbilt University, Nashville, TN, USA; Institute of Biophysics, CIPMM, School of Medicine, Saarland University, Homburg, Germany; current address: University Heart Center Frankfurt, Frankfurt, Germany; Department of Cardiology, University Hospital Leipzig, Germany; Medical Clinic 1, University Clinic Würzburg, Germany

## Abstract

Hypertrophic cardiomyopathy (HCM) is the most common inherited cardiac disease and caused by genetic variants that often increase sarcomeric Ca^2+^ sensitivity. While Ca^2+^ sensitization explains diastolic dysfunction, the genesis of ventricular arrhythmias is unresolved. Here, we show that HCM mutations or pharmacological interventions that increase myofilament Ca^2+^ sensitivity generate bioenergetic mismatch and oxidative stress during β-adrenergic stimulation which provide a trigger and a substrate for arrhythmias. For any given sarcomere shortening that produces work and consumes ATP, less Ca^2+^ stimulates the Krebs cycle to maintain mitochondrial NADH. This reverses the mitochondrial transhydrogenase to regenerate NADH from NADPH, supporting ATP production at the cost of NADPH-dependent antioxidative capacity. The ensuing overflow of reactive oxygen species (ROS) from mitochondria and glutathione oxidation induce spontaneous Ca^2+^ release from the sarcoplasmic reticulum and Ca^2+^ waves, well-defined triggers of arrhythmias. Furthermore, transhydrogenase-dependent ROS formation slows electrical conduction during β-adrenergic stimulation *in vivo*, providing a substrate for arrhythmias. Chronic treatment with a mitochondrially-targeted ROS scavenger abolishes the arrhythmic burden during β-adrenergic stimulation in HCM mice *in vivo,* while inducing mitochondrial ROS with a redox cycler is sufficient to induce arrhythmias in wild-type animals. These findings may lead to new strategies to prevent sudden cardiac death in patients with HCM.

---

Hypertrophic cardiomyopathy (HCM) is the most frequent inherited cardiac disease, affecting up to 0.6% of the population.^1^ HCM is characterized by abnormal hypertrophy of the left ventricle (LV), hypercontractility and diastolic dysfunction.^2–4^ Furthermore, it is one of the most common causes of sudden cardiac death (SCD) in young people^4^ and competitive athletes^5^ who die of ventricular arrhythmias. The majority of mutations in patients with HCM affects genes that encode sarcomeric proteins, with β-myosin heavy chain (*MYH7*, β-MHC),^6^ cardiac myosin-binding protein C (*MYBPC3*, cMyBPC)^7,8^ and cardiac troponin T (*TNNT2*, cTnT)^8^ accounting for two thirds of cases.^2^ These mutations have in common that they increase the affinity of myofilaments to calcium (Ca^2+^), causing hyperdynamic contraction and impaired relaxation.^9^ Recent genomic, proteomic, metabolomic and functional studies revealed mitochondrial dysfunction and oxidative stress in advanced stages of HCM, in association with the severity of the disease.^10–14^ However, the actual mechanisms how sarcomeric mutations can induce such defects, and how these relate to arrhythmias, is currently unresolved.^4^

Previously, we revealed that Ca^2+^ sensitization by HCM-specific mutations increases cytosolic Ca^2+^ buffering, which prolongs cardiac action potential duration and predisposes to ventricular ectopy and sustained tachycardia.^15,16^ Furthermore, HCM-causing mutations in *MYH7* or *TNNT2* increase the energetic cost of contraction and relaxation,^17,18^ inducing focal energy deprivation that further increases susceptibility towards arrhythmias through slowing conduction velocity.^19^ While these data indicate that alterations of Ca^2+^ handling and mitochondrial energetics are central to the genesis of arrhythmias, the exact interplay between these processes is unresolved.

Force generation by myofilaments consumes large amounts of ATP that are replenished by oxidative phosphorylation in mitochondria. The Krebs cycle produces NADH which donates electrons to the electron transport chain (ETC), establishing a protonmotive force across the inner mitochondrial membrane that drives ATP production at the F_1_F_o_ ATP-synthase (**Supplemental Fig. 1a**).^20,21^ When ATP consumption increases, ADP accelerates electron flux along the ETC, oxidizing NADH to NAD^+^. Physiological increases of workload are triggered by β-adrenergic stimulation, enhancing the rates and amplitudes of cytosolic Ca^2+^ transients. This promotes uptake of Ca^2+^ into mitochondria, where it stimulates Krebs cycle dehydrogenases to match NADH supply to the elevated demand. Such “parallel activation” of ATP consumption and regeneration through Ca^2+^ maintains NADH/NAD^+^ in a reduced redox state for constant electron supply to the ETC.^20,22,23^ This model predicts that any increase in myofilament Ca^2+^ sensitivity, be it by HCM mutations or pharmacological Ca^2+^ sensitization, should induce a mismatch between ATP consumption and mitochondrial Ca^2+^-stimulated regeneration and thereby cause oxidation of the NADH/NAD^+^ ratio.

Besides providing electrons for ATP production, NADH regenerates NADPH via the nicotinamide nucleotide transhydrogenase (NNT), an enzyme located in the inner mitochondrial membrane whose reaction (NADH+NADP^+^ ↔ NADPH+NAD^+^) is coupled to the proton gradient, favoring NADPH regeneration in the *forward* mode (**Supplemental Fig. 1a**).^24^ NADPH equilibrates mitochondrial glutathione (GSH) via glutathione reductase (GRX), which serves as substrate for detoxification of hydrogen peroxide (H_2_O_2_) by glutathione peroxidase (GPX).^20,25^ We previously discovered that pathological cardiac afterload *reverses* the NNT reaction to regenerate NADH from NADPH to support ATP production, but at the cost of glutathione oxidation, increasing mitochondrial reactive oxygen species (ROS) emission (**Supplemental Fig. 1b**) that causes necrosis, systolic dysfunction and premature death.^26^

Here, we show that myofilament Ca^2+^ sensitization induced by HCM-specific mutations in the *MYBPC3* or *TNNT2* genes, or by the Ca^2+^ sensitizer EMD 57033, exacerbates mechanical workload in cardiac myocytes without a proportional increase in cytosolic and/or mitochondrial Ca^2+^, which oxidizes mitochondrial NADH and via reverse-mode NNT, also NADPH during β-adrenergic stimulation. This NADPH oxidation and the ensuing mitochondrial ROS overflow trigger arrhythmias, which can be prevented by scavenging H_2_O_2_ with catalase or rebalancing the NAD(P)H/NAD(P)^+^ redox state by inhibiting mitochondrial Ca^2+^ efflux. Moreover, pretreatment with SS-31, a tetrapeptide that binds to mitochondrial cardiolipin,^27^ or the mitochondria-targeted ROS scavenger ubiquinone (Mito-Q)^28^ prevented cardiac conduction abnormalities and arrhythmias, respectively, in response to β-adrenergic stimulation *in vivo* in mice carrying an HCM-typical *TNNT2* mutation. These results reveal a key missing link in HCM pathophysiology between mutation-induced derangements of EC coupling, energy supply-and-demand mismatch, oxidative stress and susceptibility to arrhythmias. Furthermore, we provide proof of concept that clinically safe and applicable agents targeting mitochondria may prevent potentially lethal arrhythmias in patients with HCM.

## Methods

A detailed description of the Methods is available in the online supplement.

### Animal models and handling

Animal procedures were approved by the local animal ethics committee and conducted in accordance with institutional guidelines. Mice that carry the human c.772G>A *MYBPC3* mutation at the homozygous state (*Mybpc3*-KI) and WT controls were generated in a Black Swiss background as described previously^29^ and used at an age of 12-16 weeks (**Fig. 1, 2a-c**). For experiments using EMD 57033 (**Fig. 2d-i**), myocytes were isolated from 12 weeks old C57BL/6N mice that were obtained from Charles River (Sulzfeld, Germany; C57BL/6NCrl, strain code 027). Transgenic mice expressing human mutant or the WT troponin T (*Tnnt2*-I79N and -WT) were generated as described previously^30^ in a mixed genetic background and used at an age of 12-16 weeks (**Fig. 4**). For *in vivo* experiments, non-transgenic mice expressing the endogenous (murine) WT *Tnnt2* (NTG) were used as controls (**Fig. 5**, **7d-f**). Sperms of mice overexpressing catalase specifically in mitochondria (mCAT)^31^ were obtained from Peter S. Rabinovitch (University of Washington, US) and mice rederived in a C57BL/6N background (**Fig. 3**). Littermate mice without mCAT expression were used as controls. Mice with global expression of the mitochondria-located H_2_O_2_ sensor mito-roGFP2-Orp1^32^ or with cardiac myocyte-specific expression of cyto-or mito-Grx1-Orp2^33^ (**Fig. 6**) were generated previously^32,33^ and held on a C57BL/6N background, respectively.

**Figure 1:**
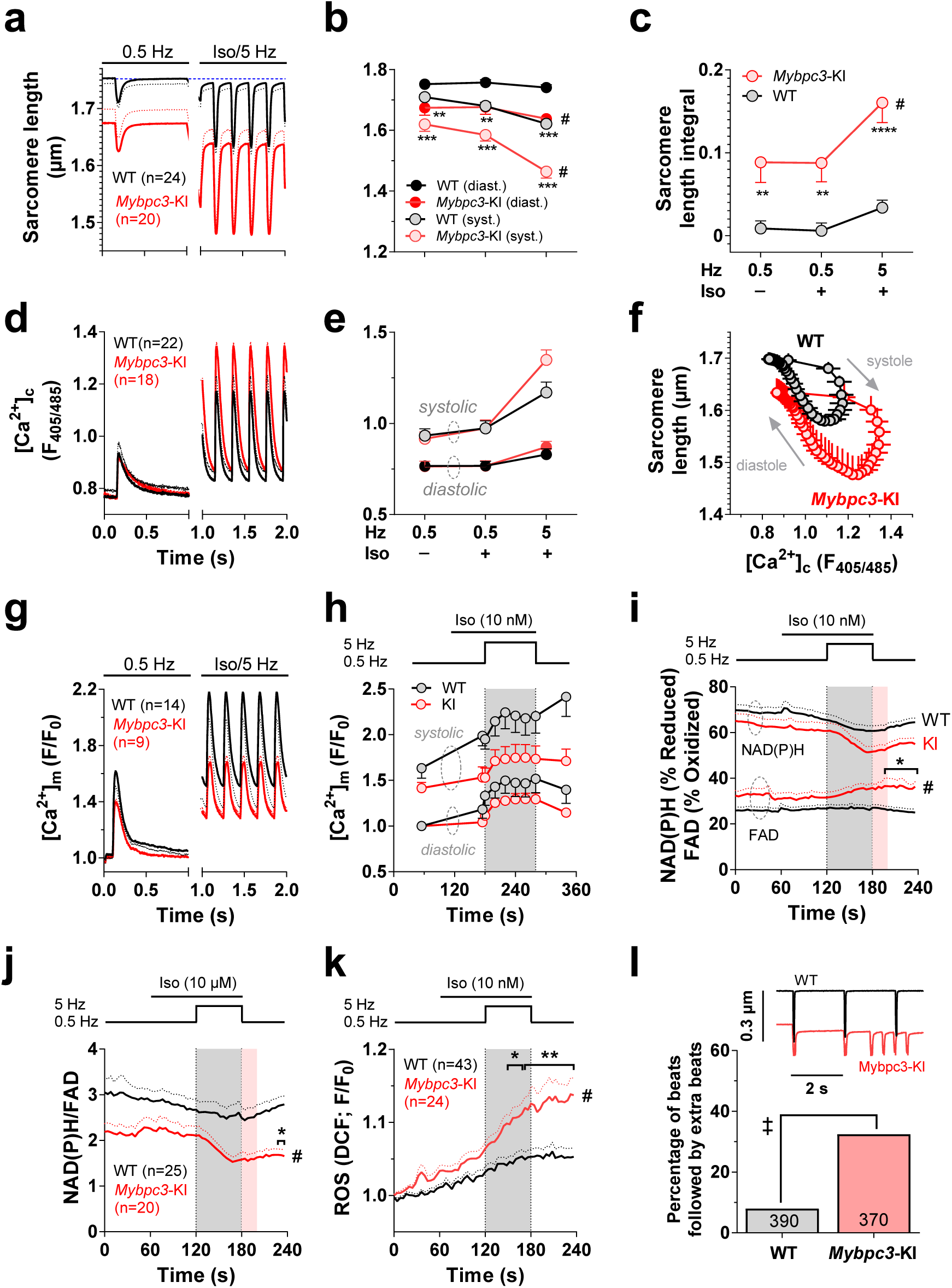
Energetic mismatch in *Mybpc3*-KI myocytes increases mitochondrial ROS emission and is associated with arrhythmias Cardiac myocytes were isolated from *Mybpc3*-KI and WT mice and electrically stimulated at 0.5 Hz. Then isoproterenol (Iso; 10 nM) was washed in and after 1 min, stimulation rate was increased to 5 Hz for 1 min. Thereafter, Iso was washed out and stimulation rate set back to 0.5 Hz. Diastolic and systolic sarcomere length were recorded in wild-type (WT; n=24 cells of n=2 mice) and *Mybpc3*-KI myocytes (KI; n=20/2) and illustrated as averaged original traces (**a**) or single data points (**b**). Under similar conditions and at similar time points, systolic and diastolic [Ca^2+^]_c_ and the amplitude of [Ca^2+^]_c_ are illustrated in **d** and **e**. **c**, Sarcomere length integral, calculated as the area under the curve between the traces and the dashed blue line in **a**. **f**, [Ca^2+^]_c_ plotted against sarcomere length at 5 Hz and Iso, obtained in the same cells, respectively. Mitochondrial Ca^2+^ ([Ca^2+^]_m_) transients (**g**) and averaged values for systolic and diastolic [Ca^2+^]_m_ (**h**) from patch-clamped *Mybpc3*-KI (n=9/7) and WT myocytes (n=14/8). **i**, Reduced NAD(P)H and oxidized FAD in *Mybpc3*-KI (n=20/2) and WT myocytes (n=25/2), respectively. **j**, Ratio of NAD(P)H/FAD in both genotypes (data from **i**). **k**, Intracellular ROS accumulation reported by DCF fluorescence in WT (n=43/6) or *Mybpc3*-KI myocytes (n=24/5). **l**, Quantification (*main figure*) and representative example (*inset*) of unstimulated extra beats within the first 20 s after stepping back from 5 to 0.5 Hz (red shaded area in **j**) in myocytes of *Mybpc3*-KI (n=37/3) and WT mice (n=39/3). Error bars indicate standard errors of the mean (SEM); n-numbers are indicated as numbers of cells/mice (**a-k**) or numbers of beats (10 per cell; **l**); *p<0.05, **p<0.01, ****p<0.0001 (Bonferroni post-test) and ^#^p<0.05 (2-way ANOVA); ^‡^p<0.0001 Chi-square test.

**Figure 2:**
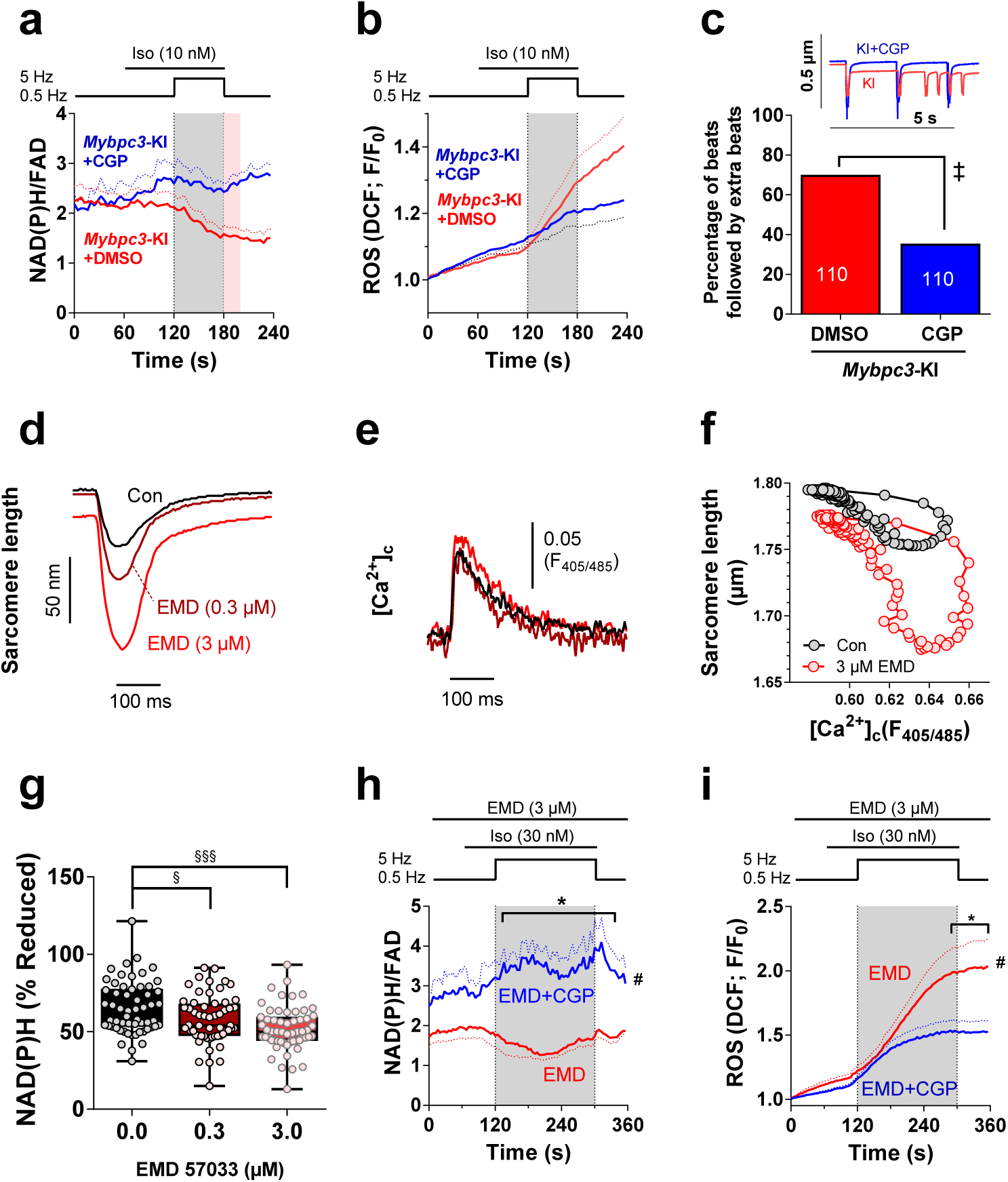
Inhibiting mitochondrial Ca^2+^ export prev s mitochondrial oxidation and ROS emission in response to myofilament Ca^2+^ sen tization. Cardiac myocytes from *Mybpc3*-KI mice were exposed to the same protocol as in Figure 1, but in the presence of an inhibitor of the mitochondrial Na^+^/Ca^2+^-exchanger, CGP-37157 (CGP, 10 µM; n=11 cells of n=4 mice) or vehicle (DMSO; n=11/4), and NAD(P)H/FAD (**a**) or ROS accumulation (DCF fluorescence; **b**; n=13/11) were determined. **c**, Representative examples (*inset*) and statistical analysis (*main figure*) of extra beats occurring in the first 20 s after reducing stimulation rate from 5 to 0.5 Hz (red shaded area in **a**). Representative sarcomere length (**d**) and [Ca^2+^]c (**e**) in a cardiac myocyte from C57BL/6N mice stimulated at 1 Hz in the absence (Con) and presence of 0.3 or 3 µM EMD 57033 (EMD), respectively. **f**, Sarcomere length (of the same cell as in **d, e**) plotted against [Ca^2+^]c in the absence or presence of 3 µM EMD, respectively. **g**, NAD(P)H/FAD in myocytes paced at 1 Hz in the absence and presence of 0.3 or 3 µM EMD, respectively (n=58/14). NAD(P)H/FAD (**h**) and ROS accumulation (DCF fluorescence; **i**) in C57BL/6N myocytes exposed to a similar stress protocol as described in Figure 1, in the absence (EMD; n=5/3 for NAD(P)H/FAD and n=12/3 for ROS) or presence of 10 µM CGP-37157 (CGP; n=4/2 & 12/2), respectively. Error bars indicate the standard errors of the mean (SEM); n-numbers are indicated as numbers of myocytes/mice (**g**) or numbers of analysed beats (10 per cell; **c**), respectively. ^+^p<0.05 unpaired t-test; *p<0.05, (Bonferroni post-test) and ^#^p<0.05 (2-way ANOVA).

**Figure 3:**
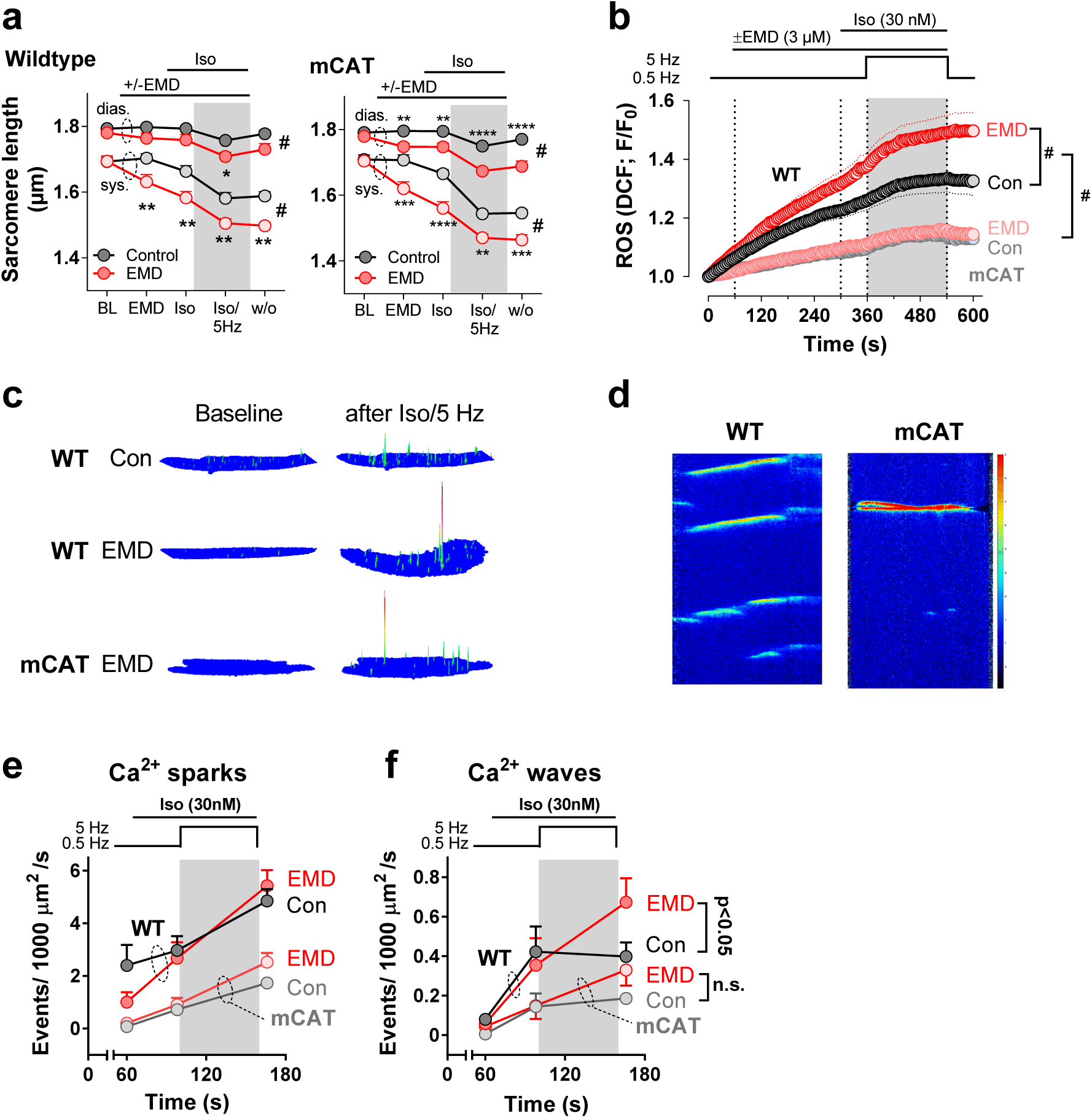
Ca^2+^ sensitization with EMD-57033 provokes mitochondrial H_2_O_2_ formation, Ca^2+^ sparks and Ca^2+^ waves Cardiac myocytes were isolated from mice overexpressing catalase specifically in mitochondria (mCAT) or wildtype littermate control mice (WT), respectively, in the presence of the Ca^2+^ sensitizer EMD-57033 (EMD; WT, n=31/10; mCAT, n=28/9) or DMSO as vehicle (Control, Con; WT, n=29/10; mCAT, n=26/9), respectively, and then exposed to isoproterenol (30 nM) and 5 Hz stimulation. Systolic and diastolic sarcomere length were recorded (**a**) together with the fluorescence of the ROS-indicator DCF (**b**). In a second set of experiments, Ca^2+^ sparks (**c, e**) and waves (**d, f**) were recorded in WT or mCAT myocytes preincubated with EMD-57033 (EMD; WT, n=54; mCAT, n=61) or vehicle (Con; WT, n=52; mCAT, n=54), respectively, during a 6.8 s unstimulated period after being exposed to 0.5 Hz stimulation in the absence (1 min) and then presence of 30 nM isoproterenol (30 s), and then after 1 min of 5 Hz stimulation in the maintained presence of isoproterenol, respectively. Error bars indicate the standard errors of the mean (SEM); n-numbers are indicated as numbers of cells/mice; *p<0.05, **p<0.01, ***p<0.001, ****p<0.0001 (Bonferroni post-test) and ^#^p<0.05 (2-way ANOVA).

**Figure 4:**
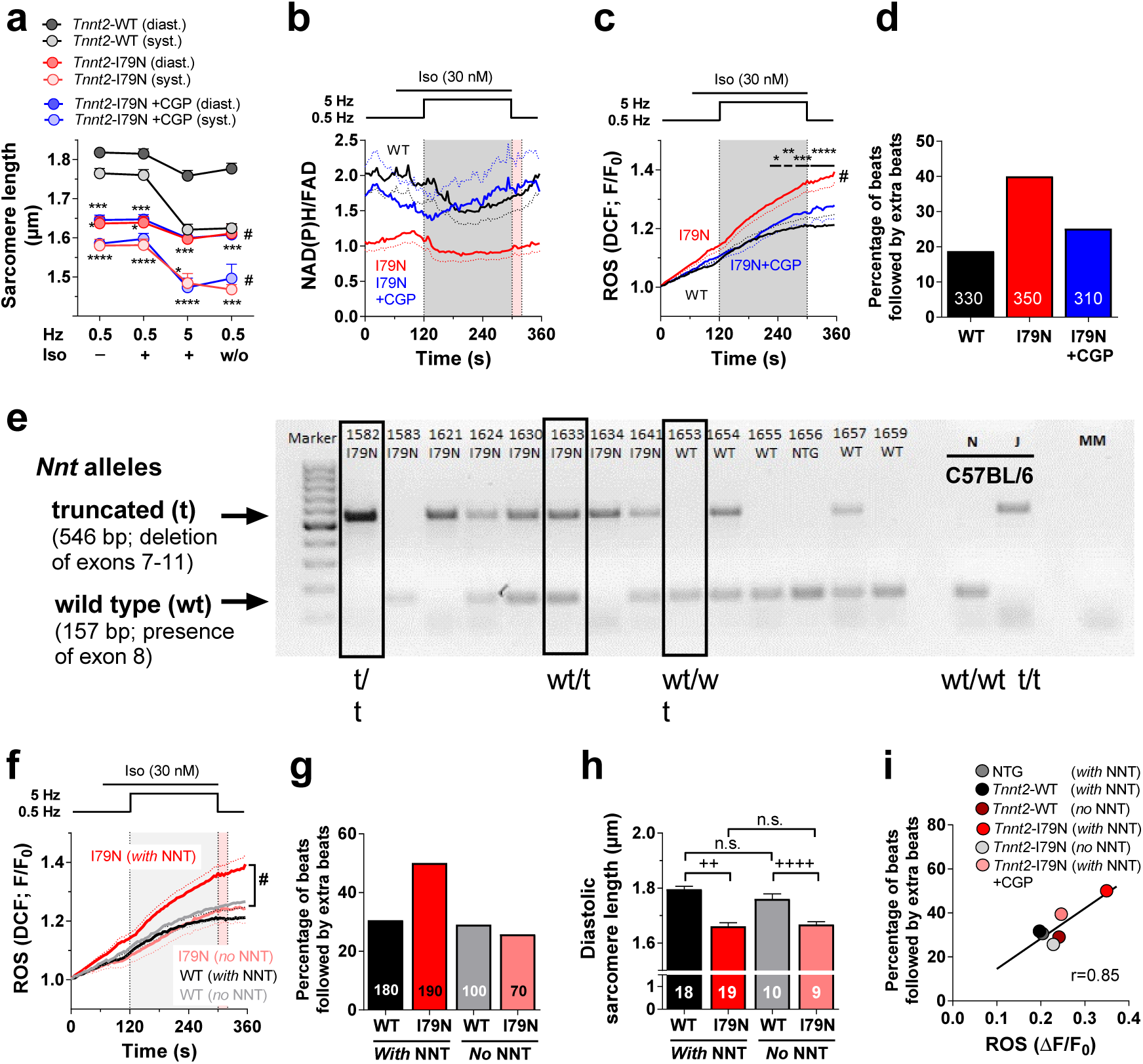
Isoproterenol-induced mitochondrial ROS emission and arrhythmias in *Tnnt2*-I79N myocytes require a functional transhydrogenase. Cardiac myocytes from transgenic mice carrying either an additional wild-type (WT) or a mutant cardiac troponin T (*Tnnt2*) cDNA (I79N) were exposed to a similar stress protocol as described in Fig. 1, with *Tnnt2*-I79N myocytes examined in the absence or presence of CGP-37157 (CGP; 10 µM). Together with sarcomere length (**a**; WT, n=32/7; I79N, n=27/6; I79N+CGP, n=26/6), NAD(P)H/FAD (**b**; WT, n=21/4; I79N, n=12/3; I79N+CGP, n=9/3) or ROS accumulation (**c**; WT, n=18/5; I79N, n=19/10; I79N+CGP, n=18/9) were recorded and after the stress protocol, extra beats were quantified (**d**; WT, n=33/7; I79N, n=35/8; I79N+CGP, n=31/8). **e**, Original PCR analysis of genotyping for *Nnt* with either truncated (t) or wild-type (wt) alleles (truncated gene, product of 546 bp; wt gene, 157 bp; see Methods). C57BL/6N (N) and C57BL/6J (J) mice served as controls with homozygous expression of wt (N) or t alleles (J), respectively. MM, master mix. **f**, ROS accumulation in WT or I79N mice expressing at least one (*with* NNT; WT, n=18; I79N, n=19) or no wild-type allele for the *Nnt* (*no* NNT; WT, n=10; I79N, n=7), respectively. **g**, Percentage of beats followed by extra beats and **h**, diastolic sarcomere length in the same genotypes as indicated in **f**. **i**, The same quantification of cellular arrhythmias as in **g** plotted against the net accumulation of ROS in the same group, respectively. Error bars or dotted lines indicate the standard errors of the mean (SEM); n-numbers are indicated as numbers of cells/mice (**f**) or numbers of analysed beats (10 per cell; **d, h**); *p<0.05, **p<0.01, ***p<0.001, ****p<0.0001 (Bonferroni post-test) and ^#^p<0.05 (2-way ANOVA); ^a^p<0.0001 Chi-square test, I79N-Nnt^wt/t^ versus all other groups; ^++^p<0.01 and ^+++^p<0.001, unpaired t-test.

**Figure 5:**
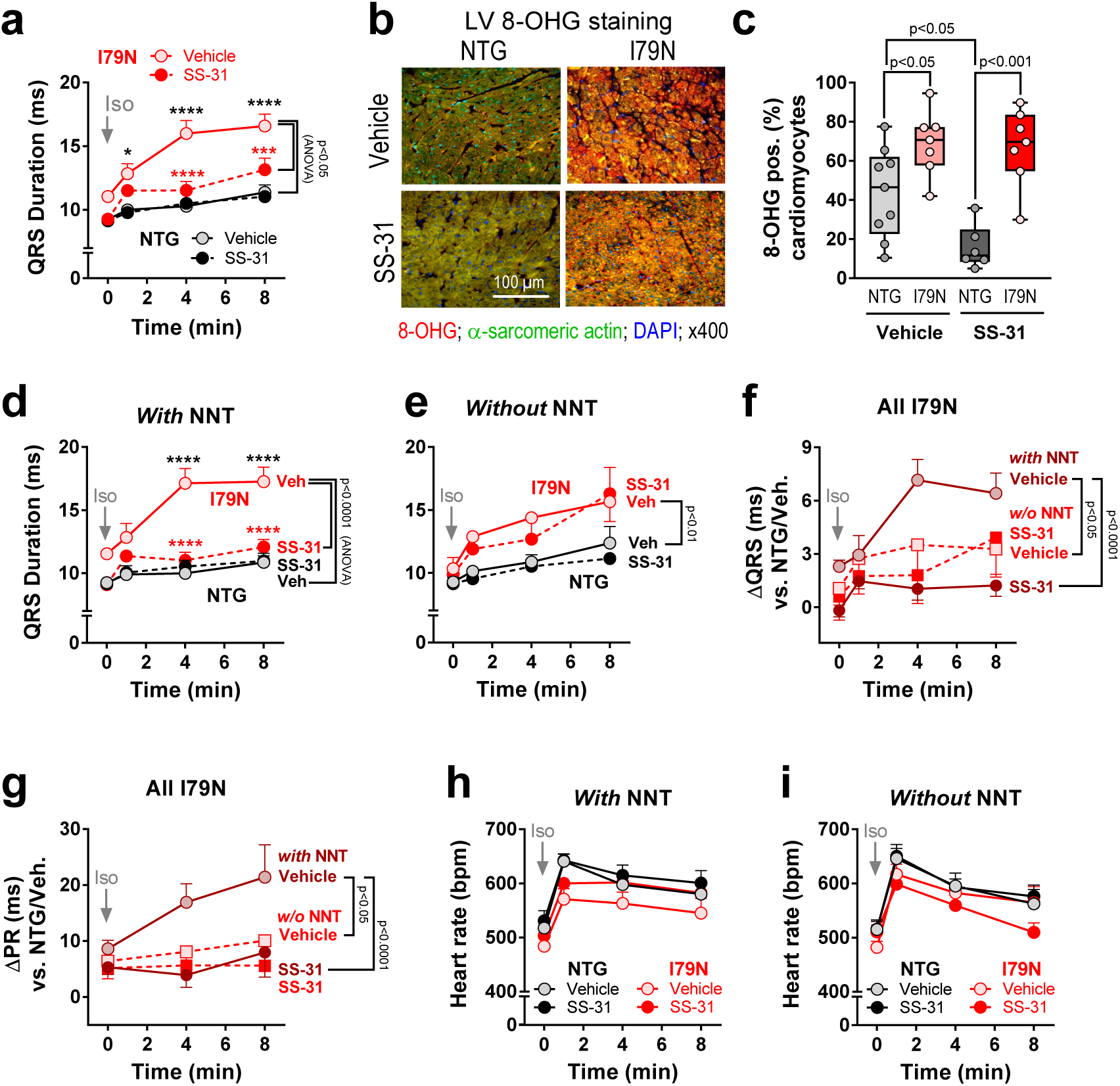
Targeting mitocho rial cardiolipin prevents QRS broadening in a transhydro fashion in *Tnnt2*-I79N hearts. Anesthetized *Tnnt2*-I79N or non-transgenic mice (NTG) received either vehicle (Veh; 0.9% NaCl; NTG, n=20; *Tnnt2*-I79N, n=17) or SS-31 (3 mg/kg; NTG, n=17; *Tnnt2*-I79N, n=14) per i.p. injection one hour prior to i.p. isoproterenol (Iso; 3 mg/kg) injection. **a**, QRS duration (ms) before and after Iso injection. Left ventricular (LV) 8-OHG staining as representative images (**b**) or quantification (**c**) in a subset of the hearts displayed in **a** (Vehicle: NTG, n=9; I79N, n=7; SS-31: NTG, n=7; I79N, n=7). **d and e**, same analysis as in **a**, but stratified by *Nnt* genotype as either all mice harboring at least one functional allele of the *Nnt* gene and thereby, with a functional NNT protein (**d;** NTG+Veh, n=13; NTG+SS-31, n=10; I79N+Veh, n=10; I79N+SS-31, n=12) or two truncated *Nnt* alleles and thereby, lack of any functional NNT protein (**e**; NTG+Veh, n=7; NTG+SS-31, n=7; I79N+Veh, n=7; I79N+SS-31, n=2), respectively. **f,** Isoproterenol-induced change in QRS duration related to the average QRS duration in vehicle-treated NTG mice at the respective time points. **g**, same as in **f**, but isoproterenol-induced change in PR intervals. **h** and **i**, isoproterenol-induced changes of heart rate in mice with either expression of at least one (**h**) or no functional Nnt allele (**i**), respectively. Error bars indicate the standard errors of the mean (SEM); n-numbers are indicated as numbers of mice/hearts. Unpaired t-Test was performed for un-matched variables. Comparison between groups was performed for vehicle and SS31-treated animals.*p<0.05, **p<0.01, ***p<0.001, ****p<0.0001 (Bonferroni post-test; black: NTG/vehicle vs. I79N/vehicle; red: I79N vehicle vs. SS-31) and #p<0.05 (2-way ANOVA).

**Figure 6:**
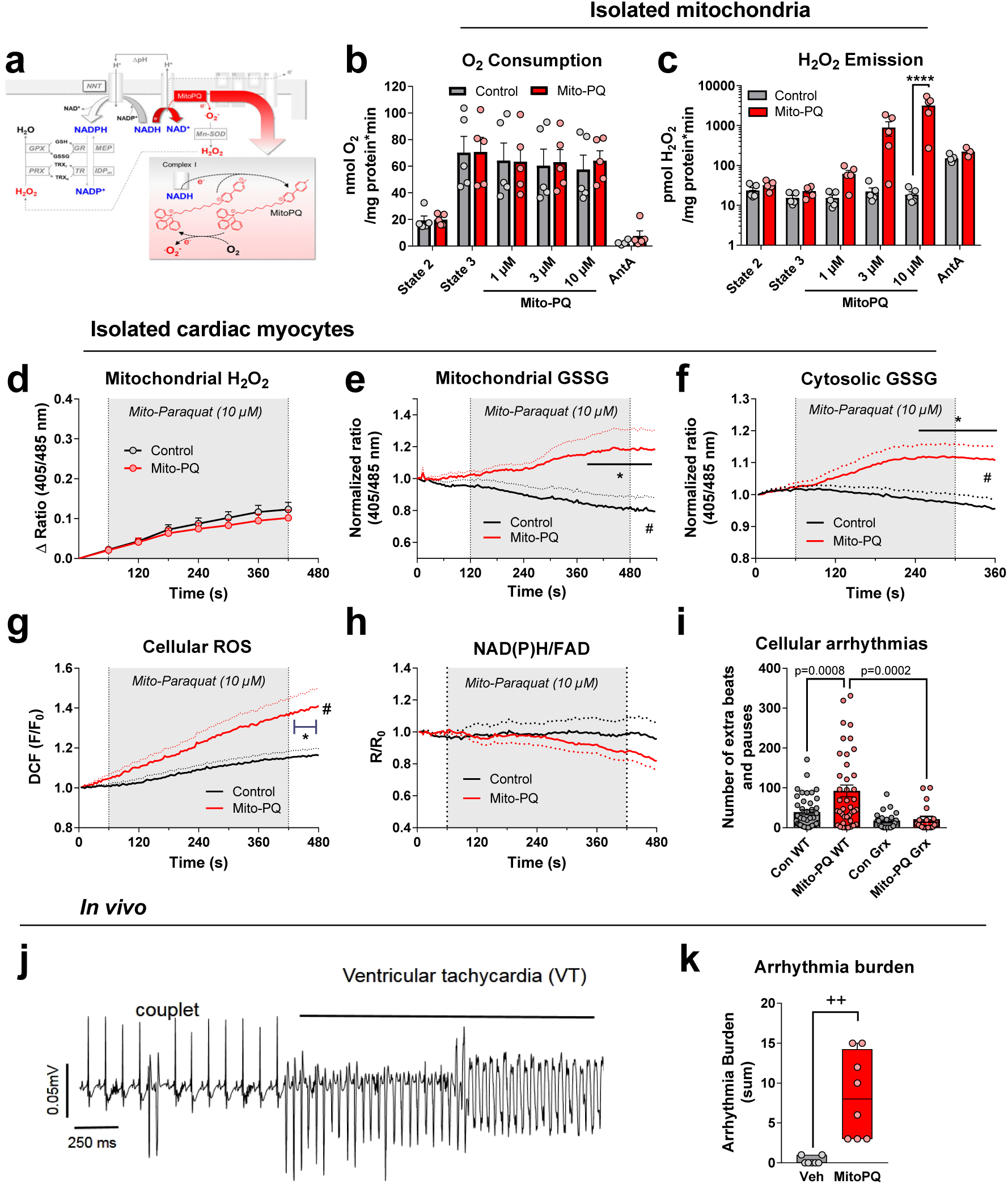
Mitochondrial glutathione oxidation induces arrhythmias **a**, Schematic illustration how mitochondrial paraquat (Mito-PQ) oxidizes NAD(P)H by transferring electrons from NADH via complex I to paraquat, which reduces oxygen (O2) to superoxide (.O2^-^). After dismutation to H2O2, NADPH-coupled enzymes eliminate H2O2. **b** and **c**, Effect of 1-10 µM Mito-PQ (n=5) or vehicle (Control; n=5) on O2 consumption (**b**) and H2O2 emission (**c**) in mitochondria supplied with pyruvate/malate (5/5 mM) respiring in state 3 (in the presence of 100 µM ADP). AntA, addition of antimycin A to block complex III, which is also a positive control for H2O2 emission. **d**-**h**, Cardiac myocytes were paced at 1 Hz and exposed to Mito-PQ (10 µM) or vehicle (Control), and the effect on mitochondrial H2O2 (**d**; indicated by the ratio of emissions of Mito-roGFP2-Orp1 at 405/485 nm; Control, n=8; Mito-PQ, n=12), oxidized glutathione (GSSG) in mitochondria (**e**; normalized ratio of emissions of Mito-Grx1-roGFP2 at 405/485 nm; Control, n=7/2; Mito-PQ, n=6/2) or the cytosol (**f**; normalized ratio of Cyto-Grx1-roGFP2 fluorescence at 405/485 nm; Control, n=23/4; Mito-PQ, n=18/4), the fluorescence of the non-specific cellular ROS indicator DCF (**g**; Control, n=10; Mito-PQ, n=11) and the normalized ratio of the autofluorescence of reduced NAD(P)H divided by oxidized FAD (**h**; Control, n=36/8; Mito-PQ, n=38/8) were detected. **i**, Number of extra beats and pauses induced by Mito-PQ or vehicle in isolated cardiac myocytes without any fluorescence reporter (Vehicle, n=39; Mito-PQ, n=40), or in cardiac myocytes expressing Cyto-Grx1-roGFP2 (Vehicle, n=23; Mito-PQ, n=20) were quantified. Original recording (**j**) and quantification of the arrhythmic burden (**k**; a score quantifying PVCs, AV blocks and VTs) in NTG mice pretreated with either Mito-PQ (0.1 mg/kg i.p.; n=8) or vehicle (n=7), respectively, and then injected with isoproterenol (1.5 mg/kg i.p.). Error bars indicate the standard errors of the mean (SEM); n-numbers are indicated as numbers of cells and number of animals (**d-i**) or mice (**b,c,k**); *p<0.05, ****p<0.001 (Bonferroni post-test) and ^#^p<0.05 (2-way ANOVA); ^+^p<0.05 and ^++^p<0.01, unpaired t-test.

### Statistical analyses

Values are displayed as means ±SEM unless otherwise indicated. Two-way ANOVA followed by Bonferroni multiple comparisons test, paired and unpaired Student’s t-tests, Fisheŕs exact test and Kaplan-Meier survival analyses were performed using GraphPad Prism version 6.00 for Windows, GraphPad Software, La Jolla California USA, www.graphpad.com. Nested t-test or 1-way ANOVA were performed using GraphPad Prism version 10 for Windows.

GraphPad Prism was used for graph generation and statistical evaluation. Groups were evaluated as non-parametric with Mann-Whitney for 2 groups and Kruskal-Wallis with Dunn’s correction for multiple comparisons.

For cellular experiments, we performed an *a priori* power calculation using G*Power according to Faul et al.^34^ that was conducted for two independent groups (WT/*Mybpc3*-KI), considering sarcomere length as a critical parameter for diastolic dysfunction as a readout. For calculating the estimated effect size, we relied on our previous experiments reported in Flenner et al.,^35^ in which sarcomere length was 1.82 ± 0.05 µm in WT and 1.76 ± 0.05 µm in KI mice, resulting in an effect size of 1.20. Alpha and beta error probability were set at 0.05, respectively. We aimed at analyzing equally sized groups. The calculated needed sample size for an unpaired t-test was n=20 for each group. The estimates were extended to the cellular measurements of other parameters. Cells that died during the protocols or could not be calibrated were excluded from analysis.

For animal studies, we also used G*Power according to Faul et al.^34^ and considered the QRS width after isoproterenol challenge as the critical parameter for arrhythmia susceptibility based on our previous data reported by Huke et al.,^19^ where QRS width was 9.87 ± 1.48 [mean ±SD] in *Tnnt2*-WT and 12.25 ± 2.00 in *Tnnt2*-I79N mice, respectively. To detect a 60% reduction of QRS width induced by SS-31 in *Tnnt2*-I79N mice, and considering an alpha of 0.05 and a beta of 0.2, we calculated an effect size of 0.72 and a resulting n-number of 14 for paired t-test analysis (cross-over design). Animal that died after the first protocol and could not enter the second protocol were excluded from the paired analysis.

For cellular experiments, blinding was not feasible since the shorter sarcomere length revealed the HCM-typical phenotype in *Tnnt2*-I79N and *Mybpc3*-KI myocytes or in myocytes treated with EMD 57033. Investigators were blinded to the genotype of the *Nnt* in *Tnnt2*-I79N and – WT mice, since this genotype was revealed after the experiments were performed. Experiments on isolated hearts and *in vivo* were executed in an unblinded fashion.

In order to control for the variance in the population, i.e. the variance between different animals on different days of experiments, we systematically applied nested analyses (t-test or 1-way ANOVA) to all data derived from cardiac myocytes. Only if *i)* the nested analysis identified a significant clustering of results from different days of experiments (i.e. of cells from different mice on different days), AND *ii)* if the nested analysis obtained a p-value >0.05 for the actual analysis, we discarded the significance obtained by our prior (non-nested) test of this data point. Otherwise, we maintained (and indicate) the results of the original analyses, since we considered a 2-way ANOVA (where appropriate) as being more adequate for multiple comparison than a (nested) t-test.

The datasets generated during and/or analysed during the current study are available from the corresponding author on reasonable request.

## Results

### HCM-specific MYBPC3 mutation induces energetic mismatch in mitochondria

To elucidate the impact of mutations of sarcomeric proteins on mitochondrial function and redox state, we used a *Mybpc3*-targeted knock-in mouse model of HCM, which carries the human c.772G>A *MYBPC3* mutation at the homozygous state (*Mybpc3*-KI).^29^ These mice exhibit increased myofilament Ca^2+^ sensitivity and diastolic dysfunction.^35,36^ Diastolic sarcomere length of isolated cardiac myocytes was shorter in *Mybpc3*-KI than in wild-type (WT) cells at 0.5 Hz despite no difference in diastolic cytosolic Ca^2+^ concentrations ([Ca^2+^]_c_; **Fig. 1a, b, d, e**). While the rate of sarcomere re-lengthening was similar, the rate of [Ca^2+^]_c_ decay was longer in *Mybpc3*-KI than in WT myocytes (**Supplemental Fig. 2a**, **b**). Fractional sarcomere shortening and the amplitudes of [Ca^2+^]_c_ transients did not differ between genotypes at 0.5 Hz (**Supplemental Fig. 2a**, **c**).

Since in young patients with HCM, arrhythmias often occur during physical activity,^37–39^ we simulated physiological workload by exposing myocytes to the β-adrenergic receptor agonist isoproterenol (10 nM) and elevated stimulation rate to 5 Hz for 60 seconds.^26,35,36^ During this workload transition, diastolic and systolic sarcomere length shortened even further in *Mybpc3*-KI versus WT myocytes (**Fig. 1a, b**). Furthermore, fractional sarcomere shortening increased more in *Mybpc3*-KI than in WT myocytes, confirming our previous data^35^ (**Extended data, Fig. 2c**; *upper trace*). In contrast, no differences in systolic and diastolic [Ca^2+^]_c_, amplitude of [Ca^2+^]_c_ transients, nor the kinetics of sarcomere re-lengthening and [Ca^2+^]_c_ decay were observed between genotypes during β-adrenergic stimulation at 5 Hz (**Fig. 1d, e**; **Supplemental Fig. 2b**, **c**).

Considering that *i)* sarcomere length of unloaded myocytes predicts elevated tension under loaded conditions,^40^ and *ii)* the energetic cost of diastolic tension is comparable to that of systolic force development,^41^ we calculated the sarcomere length integral as an index for the sum of systolic and diastolic mechanical work performed by myocytes. This integral was substantially higher in *Mybpc3*-KI than in WT myocytes (**Fig. 1c**). When plotting sarcomere length against the respective [Ca^2+^]_c_ at 5 Hz in the presence of β-adrenergic stimulation, sarcomere length was shorter at any given [Ca^2+^]_c_ during systole and diastole in *Mybpc3*-KI myocytes (**Fig. 1f**). These data indicate that elevated myofilament Ca^2+^ affinity increases the mechanical work of a myocyte at any given [Ca^2+^]_c_ in *Mybpc3*-KI mice.

Elevated systolic or diastolic tension both increase O_2_ consumption,^41^ since ADP (the product of ATP hydrolysis) accelerates electron flux at the respiratory chain (**Supplemental Fig. 1a**).^23^ This oxidizes NADH, which is matched by Ca^2+^-induced acceleration of Krebs cycle activity under physiological conditions.^23^ Since increased myofilament Ca^2+^ sensitivity may additionally buffer [Ca^2+^]_c_,^16^ and the efficacy of mitochondrial Ca^2+^ uptake depends on the rate of rise of [Ca^2+^]_c_,^42^ we determined mitochondrial and cytosolic Ca^2+^ by a patch-clamp based approach.^26,36,43,44^ Despite a trend towards lower beat-to-beat transients and less accumulation of mitochondrial Ca^2+^ ([Ca^2+^]_m_) in *Mybpc3*-KI myocytes, these were not significantly different from WT (**Fig. 1g, h**). In any case, the elevated mechanical workload in *Mybpc3*-KI myocytes (**Fig. 1c**) was not adequately matched by congruently increased mitochondrial Ca^2+^ uptake and could therefore provoke an energetic imbalance.

### Energetic mismatch oxidizes pyridine nucleotides in Mybpc3-KI cardiomyocytes

The redox states of NADH/NAD^+^ and FADH_2_/FAD are under the control of ADP and Ca^2+^, with Ca^2+^ *reducing* the redox states through Krebs cycle activation, and ADP *oxidizing* them through acceleration of respiration (**Supplemental Fig. 1a**).^23^ In agreement with our hypothesis and the observed mismatch between workload and [Ca^2+^]_m_, NAD(P)H/NAD(P)^+^ and FADH_2_/FAD (assessed by their autofluorescence) were more oxidized in *Mybpc3*-KI than in WT myocytes already at low stimulation rates, which was further aggravated during 5 Hz and β-adrenergic stimulation (**Fig. 1i**). Accordingly, the ratio of NAD(P)H/FAD (as a sensitive index of the mitochondrial redox state that eliminates movement artifacts^26,36,44^) was more oxidized in *Mybpc3*-KI versus WT at baseline and, in particular, during the physiological stress protocol (**Fig. 1j**). Hence, the mismatch between elevated mechanical load and mitochondrial Ca^2+^ oxidizes the pyridine nucleotides NADH, FADH_2_ and NADPH in *Mybpc3*-KI myocytes.

### Energetic mismatch increases mitochondrial ROS in Mybpc3-KI cardiomyocytes

Since H_2_O_2_-eliminating enzymes require NADPH, and NADPH is regenerated by reactions that rely on NADH and other products of the Krebs cycle (**Supplemental Fig. 1a**),^45^ decreased mitochondrial Ca^2+^ uptake induces NADPH oxidation and increased mitochondrial H_2_O_2_ emission.^46^ Accordingly, emission of H_2_O_2_ (reported by the non-specific ROS-indicator DCF, together with other ROS) was higher during the workload transition in *Mybpc3*-KI than in WT myocytes (**Fig. 1k**). Plotting NAD(P)H/FAD against ROS accumulation further illustrates that mitochondrial pyridine nucleotide oxidation was closely associated with ROS emission (**Supplemental Fig. 2d**).

To rule out that elevated ROS emission was related to defects of the ETC and/or the anti-oxidative capacity *per se*, we isolated cardiac mitochondria from *Mybpc3*-KI and WT hearts and determined respiration, superoxide (.O_2_) formation and H_2_O_2_ emission in the absence and presence of ADP, and found no differences between genotypes for these parameters (**Supplemental Fig. 3**). Accordingly, mitochondrial .O_2_ formation and membrane potential (ΔΨ_m_), measured in intact cardiac myocytes during the stress protocol, did not differ between *Mybpc3*-KI and WT myocytes (**Supplemental Fig. 2e**, **f**). Therefore, increased mitochondrial ROS emission in *Mybpc3*-KI myocytes is not the result of defective mitochondria or increased ROS production at the ETC *per se*, but rather of the redox imbalance induced by the mismatch of regulating cellular factors (i.e., Ca^2+^ and ADP) that limits elimination of H_2_O_2_ in the matrix.

We realized that *Mybpc3*-KI myocytes were particularly sensitive to the stress protocol, with many cells displaying arrhythmias and hypercontracture (**Supplemental Fig. 2g**).^35^ Therefore, the time of 5-Hz stimulation was limited to 1 min in this model (*Mybpc3*-KI and WT), and at the end of the protocol, after stepping back to 0.5 Hz, we determined the rate of spontaneous extra contractions of myocytes (red area in **Fig. 1i, j**). We refer to these as “cellular arrhythmias”, which were more frequent in *Mybpc3*-KI than in WT myocytes (**Fig. 1l**), suggesting that mitochondrial oxidation and ROS emission may trigger these.

### Inhibiting mitochondrial Ca^2+^ export prevents mitochondrial oxidation and arrhythmias caused by myofilament Ca^2+^ sensitization

To evaluate whether oxidation of mitochondrial pyridine nucleotides (**Fig. 1i, j**) was indeed the consequence of an imbalance between mitochondrial ADP and Ca^2+^, we inhibited the mitochondrial Na^+^/Ca^2+^-exchanger (NCLX) with CGP-37157 (CGP). In previous studies, CGP elevated [Ca^2+^]_m_ and prevented NAD(P)H oxidation during workload transitions in cardiac myocytes.^43,44,47^ Here, CGP ameliorated NAD(P)H/FAD oxidation, mitochondrial ROS emission and arrhythmias in *Mybpc3*-KI myocytes (**Fig. 2a-c** and **Supplemental Fig. 4a**, **b**), while sarcomere shortening remained unaffected (not shown). These data confirm that imbalance between workload and mitochondrial Ca^2+^ accounts for pyridine nucleotide oxidation, ROS emission and arrhythmias in *Mybpc3*-KI myocytes.

If our energetic mismatch hypothesis is correct, pharmacological myofilament Ca^2+^ sensitization should induce similar abnormalities as the mutation of *Mybpc3*. Indeed, the Ca^2+^ sensitizer EMD 57033 (EMD) decreased diastolic sarcomere length and increased systolic sarcomere shortening in normal C57BL/6N myocytes, while [Ca^2+^]_c_ remained unchanged (**Fig. 2d-f** and **Supplemental Fig. 4c-e**). This led to concentration-dependent NAD(P)H oxidation at 1 Hz stimulation (**Fig. 2g**). In the presence of EMD, mitochondrial NAD(P)H/FAD oxidation and ROS emission in response to isoproterenol/5 Hz were blunted by CGP (**Fig. 2h, i**), supporting that Ca^2+^ sensitization *per se* (rather than secondary changes in *Mybpc3*-KI myocytes) accounted for mitochondrial oxidation and ROS emission.

### Mitochondrial H_2_O_2_ triggers sarcoplasmic reticulum Ca^2+^ release and waves upon Ca^2+^ sensitization

Although DCF locates primarily (but not exclusively) to mitochondria,^48^ it is a rather nonspecific ROS-indicator,^49^ and the increase in cellular DCF fluorescence cannot discriminate between different subcellular ROS sources and redox species. Therefore, we performed experiments in transgenic mice overexpressing catalase specifically in mitochondria (mCAT)^31^ and littermate control animals (WT). EMD had similar effects on diastolic and systolic sarcomere length in both genotypes (**Fig. 3a**). DCF oxidation during the stress protocol and in particular, the EMD-induced potentiation of ROS were blunted in myocytes from mCAT versus WT mice (**Figure 3b**). These data suggest that *i)* H_2_O_2_ rather than .O_2_^-^ is the reactive oxygen species accounting for the increase in DCF fluorescence and *ii)* mitochondria are the source of ROS.

Type 2 ryanodine receptors (RyR2) of the sarcoplasmic reticulum (SR) have a high number of cysteine residues that make them sensitive to redox-dependent regulation of gating.^50–53^ In fact, low levels of mitochondrial ROS provoke spontaneous SR Ca^2+^ release events (so-called Ca^2+^ sparks) in cardiac myocytes that may trigger cardiac arrhythmias.^51,54^ Therefore, we determined Ca^2+^ sparks and waves in mCAT and WT myocytes exposed to EMD and the isoproterenol/5 Hz stress protocol. At 0.5 Hz, EMD reduced Ca^2+^ spark frequency, likely explained by EMD-induced binding of free cytosolic Ca^2+^ to myofilaments, thereby reducing Ca^2+^-induced activation of RyR2 (**Fig. 3c, e**). After the addition of isoproterenol and in particular, the 5 Hz stimulation protocol, Ca^2+^ spark frequency increased substantially in WT myocytes treated with or without EMD, with a greater net increase in EMD-treated myocytes (**Fig. 3c, e**). Furthermore, the occurrence of Ca^2+^ waves across the whole cell, which corresponds with cellular contractions detected as cellular arrhythmias in our previous experiments, increased to a greater extent in EMD-pretreated cells compared to vehicle-treated cells (**Fig. 3d, f**). Under baseline conditions, but in particular in response to EMD, isoproterenol and pacing, the increases in Ca^2+^ sparks and waves were substantially suppressed in mCAT versus WT myocytes (**Fig. 3c-f**). These data indicate that in response to β-adrenergic stimulation and Ca^2+^ sensitization, spontaneous SR Ca^2+^ release events and Ca^2+^ waves can be prevented by suppressing mitochondrial H_2_O_2_.

### HCM-causing troponin T variant also causes mitochondrial oxidation and arrhythmias

To interrogate whether the observed bioenergetic deteriorations were specific for the *Mybpc3*-KI mice or rather universal also for other HCM-causing, Ca^2+^ sensitizing mutations, we performed experiments in myocytes of transgenic mice that express the human wild-type (*Tnnt2*-WT) or I79N *TNNT2* variant (*Tnnt2*-I79N) and exhibit high susceptibility towards ventricular arrhythmias during β-adrenergic stimulation.^15,16,19,55^ Similar to *Mybpc3*-KI, *Tnnt2*-I79N myocytes displayed shorter diastolic sarcomere length, but comparable fractional sarcomere shortening, pronounced NAD(P)H/FAD oxidation and ROS emission during the workload transition compared to *Tnnt2*-WT (**Fig. 4a-c**). Again, oxidation and ROS emission were associated with cellular arrhythmias (**Fig. 4d**) and prevented by CGP, while CGP did not affect systolic or diastolic tension (**Fig. 4a-d**).

### Mitochondrial transhydrogenase mediates ROS emission and arrhythmias in Tnnt2-I79N myocytes

We recently revealed that elevated cardiac afterload triggers mitochondrial ROS emission through reversal of the transhydrogenase (NNT; **Supplemental Fig. 1b**),^26^ and C57BL/6J mice lack a functional NNT.^56^ The *Tnnt2*-I79N/-WT mice used herein were on a mixed genetic background, and our initial results obtained in these mice (ignoring the genetic background) were inconclusive (not shown). Therefore, we genotyped all mice and found a mosaic pattern of *Nnt* alleles: Some mice were homozygotes for the WT allele (wt/wt), but most of them were either heterozygotes (wt/t) or homozygotes (t/t) for the truncated *Nnt* allele (**Fig. 4e**), stemming from C57BL/6J. In fact, when regrouping mice and cells according to the absence (t/t) or presence of at least one wild-type *Nnt* allele (wt/t), we found that the absence of any functional *Nnt* allele completely abrogated the increase in ROS emission (**Fig. 4f**) and cellular arrhythmias in *Tnnt2*-I79N myocytes (**Fig. 4g**), without ameliorating diastolic dysfunction (**Fig. 4h**). ROS emission and cellular arrhythmias closely correlated across all tested groups, including also non-transgenic (NTG) mice (expressing murine *Tnnt2*) from the same cohort (**Fig. 4i**). These data imply that increased ROS emission and arrhythmias in *Tnnt2*-I79N mice result from increased myofilament Ca^2+^ sensitivity, but require an intact NNT that can reverse^26^ to deplete NADPH upon ADP-induced NADH consumption (**Supplemental Fig. 1b**).

### Targeting mitochondrial cardiolipin with SS-31 prevents conduction defects in HCM during β-adrenergic stimulation

To elucidate whether mitochondrial ROS cause arrhythmias also *in vivo*, we injected isoproterenol (Iso; 1.5 mg/kg^15,16^) in *Tnnt2*-I79N or littermate NTG mice. We previously reported that Iso-induced QRS widening in *Tnnt2*-I79N mice is caused by reduced transverse conduction velocity as a result of focal ATP depletion with consequent connexin 43 dephosphorylation, generating a substrate for ventricular arrhythmias.^19^ In line with our previous data,^19^ Iso substantially prolonged the QRS interval in *Tnnt2*-I79N, but not NTG mice (**Fig. 5a**). SS-31 is a tetrapeptide that accumulates in mitochondria where it binds to cardiolipin, a key phospholipid of the inner mitochondrial membrane that governs the assembly of the respiratory chain complexes.^27^ Through its binding, SS-31 prevents cardiolipin oxidation and thereby, preserves respiratory chain function and ATP production.^27^ In *Tnnt2*-I79N mice, pretreatment with SS-31 abrogated QRS prolongation, but had no effect in NTG mice (**Fig. 5a**).

To gain additional insight into the role of mitochondrial ROS for slowing conduction in HCM, we analyzed oxidative stress by histological detection of 8-hydroxyguanosine (8-OHG) staining, which detects oxidized nucleic acids and lipids, in LV myocardium of *Tnnt2*-I79N and NTG mice treated with either vehicle or SS-31, respectively. In fact, *Tnnt2*-I79N hearts exhibited higher 8-OHG staining than NTG, but SS-31 did not prevent oxidation (**Fig. 5b, c**). This is in line with a lack of direct scavenging of either .O_2_ or H_2_O_2_ by SS-31 (**Supplemental Fig. 5**), respectively. Together, these and previous data^27,57^ indicate that conduction deficits in *Tnnt2*-I79N hearts during β-adrenergic stimulation are prevented by SS-31, likely through protecting mitochondria from ROS-induced cardiolipin damage with subsequent energetic deficit.

### Mitochondrial transhydrogenase deficiency ameliorates conduction deficits in Tnnt2-I79N mice and abolishes the protective effect of SS-31

In isolated cardiac myocytes from *Tnnt2*-WT and -I79N mice, the lack of a functional NNT prevented the Iso-induced increase in mitochondrial ROS and arrhythmias (**Fig. 4e-g**). Genotyping mice treated with Iso *in vivo* (**Fig. 5a, c**) revealed that the Iso-induced prolongation of the QRS complex, but also the PR-interval (reporting atrio-ventricular conduction) was more pronounced in *Tnnt2*-I79N hearts with at least one functional *Nnt* allele compared to hearts with two truncated *Nnt* alleles (**Fig. 5d-g**), despite similar effects of Iso on heart rate (**Fig. 5h,i**). These data suggest that also under *in vivo* conditions, (reverse mode) NNT-mediated NADPH oxidation and ROS emission aggravate the slowing of cardiac conduction. Moreover, SS-31 prevented QRS prolongation only in *Tnnt2*-I79N hearts with a functional NNT, but had no effect in the absence of a functional NNT (**Fig. 5d-g**). Together, these data indicate that SS-31 acts downstream of mitochondrial ROS, presumably by protecting mitochondria from ROS-induced damage of the ETC^27^ and a consequent energetic deficit that accounts for slowed cardiac conduction.

### Glutathione oxidation triggers ventricular arrhythmias

To further understand the link between NADPH oxidation, the ensuing oxidation of the mitochondrial and cytosolic glutathione pools, ROS and cellular arrhythmias, we evaluated the effects of the mitochondria-targeted redox cycler Mito-Paraquat (Mito-PQ), which uses NADH-derived electrons from complex I to transfer them to O_2_, forming .O_2_^-^ (**Fig. 6a**).^58^ In isolated cardiac mitochondria respiring in state 3 (with 100 µM ADP), 10 µM Mito-PQ increased H_2_O_2_ emission ∼100-fold, without affecting the rate of respiration (**Fig. 6b,c**) or the redox state of NAD(P)H/NAD(P)^+^ (**Supplemental Fig. 6a,,b**). Exposure of isolated cardiac myocytes from BL/6N mice to the same concentration of Mito-PQ increased cellular arrhythmias (**Fig. 6i**), without altering diastolic sarcomere length or systolic shortening (**Supplemental Fig. 6c,,d**). However, in isolated cardiac myocytes expressing roGFP2-Orp1, a fluorescence reporter for H_2_O_2_ localized to mitochondria (mito-roGFP2-Orp1)^32^ which is oxidized by application of 10-50 µM extracellular H_2_O_2_ (**Supplemental Fig. 6d**), Mito-PQ did not increase mitochondrial (free) H_2_O_2_ (**Fig. 6d**), despite inducing arrhythmias (**Supplemental Fig. 6e**). Conversely, in myocytes of mice expressing roGFP2 coupled to glutathione reductase 1 (Grx1) in either mitochondria (Mito-Grx1-roGFP2) or the cytosol (Cyto-Grx1-roGFP2),^33^ Mito-PQ oxidized glutathione (GSSG) in both compartments (**Fig. 6e,f**). Accordingly, also oxidation of the (rather unspecific) whole-cellular ROS reporter DCF was observed (**Fig. 6g**). In contrast, the redox state of NAD(P)H/FAD in non-transgenic BL/6N cardiomyocytes was only modestly (but insignificantly) oxidized (**Figure 6h**), in agreement with the result in isolated mitochondria (**Supplemental Fig. 6a,,b**). This lack of an effect on NAD(P)H/FAD may be related to the fact that NADPH (regenerating GSSG via GRX) contributes only ∼23%, but NADH 77% to the overall fluorescence of total NAD(P)H in cardiac mitochondria,^26^ and Mito-PQ did not affect mitochondrial respiration (**Fig. 6b**) and therefore, NADH/NAD^+^ or FADH_2_/FAD, which dominate the NAD(P)H/FAD ratio, are unaltered. Interestingly, in myocytes expressing Cyto-Grx1-roGFP2, the Mito-PQ-induced increase in cellular arrhythmias was prevented despite glutathione oxidation in the same cells, possibly related to increased H_2_O_2_ eliminating capacity by elevated expression of the Grx-coupled reporter system (**Supplemental Fig. 6f**).

We next tested the effect of Mito-PQ *in vivo* using NTG mice expressing at least one functional allele of the *Nnt*. To facilitate the detection of arrhythmias, mice were anesthetized using ketamine and xylazine, since isoflurane anesthesia can mask an arrhythmia phenotype in mice.^59,60^ Exposure to a low dose of Mito-PQ (0.1 mg/kg) had no effect on baseline ECG parameters and did not cause ventricular arrhythmias (**Supplemental Table 2**). Additional challenge with Iso (1.5 mg/kg) of mice pretreated with low-dose Mito-PQ resulted in QRS widening and ventricular arrhythmias (**Fig. 6j-k**), an ECG phenotype akin to the one of *Tnnt2*-I79N mice under similar anesthesia.^15,16^ Injection of a higher dose of Mito-PQ (0.5 mg/kg) caused QRS widening and ventricular arrhythmias even in the absence of Iso-challenge (**Supplemental Table 2**).

To analyze mitochondrial H_2_O_2_ in response to Mito-PQ *in vivo*, we employed the MitoB/MitoP technique.^61–63^ However, no increase of free H_2_O_2_ could be detected in response to 0.5 mg/kg Mito-PQ *in vivo* (**Figure 6l**) in agreement with the results with roGFP2-Orp1 in isolated cardiac myocytes (**Figure 6d**). Accordingly, no increase in free H_2_O_2_ was detected in *Tnnt2*-I79N mice compared to NTG mice, with or without additional Iso injection (**Figure 6l**).

Taken together, these data indicate that a specific increase of mitochondrial .O_2_ (with Mito-PQ) increases mitochondrial H_2_O_2_, which is rapidly buffered by mitochondrial and cytosolic glutathione-coupled GPX at the expense of glutathione oxidation. While this glutathione oxidation in the mitochondria and cytosol is sufficient to induce arrhythmias, an increase in “free” mitochondrial H_2_O_2_ is prevented by the glutathione-based buffering systems at the subcellular site of the proposed H_2_O_2_ effect on RyR2 (**Supplemental Fig. 6f**).

### Mitochondrially-targeted therapies prevent ventricular arrhythmias in HCM

Finally, we tested the therapeutic relevance of this newly identified mechanism for arrhythmias in HCM. High workload by rapid pacing of isolated Langendorff-perfused hearts induced NAD(P)H oxidation in *Tnnt2*-I79N, but not NTG hearts (**Fig. 7a**). This NAD(P)H oxidation was accompanied by frequent episodes of non-sustained ventricular tachycardia in *Tnnt2*-I79N hearts (**Fig. 7b-c**). Analogous to its effects in isolated cardiomyocytes (**Fig. 4b,d**), pretreatment with the NCLX inhibitor CGP completely abrogated NAD(P)H oxidation and arrhythmias in *Tnnt2*-I79N hearts (**Fig. 7b-c**).

**Figure 7:**
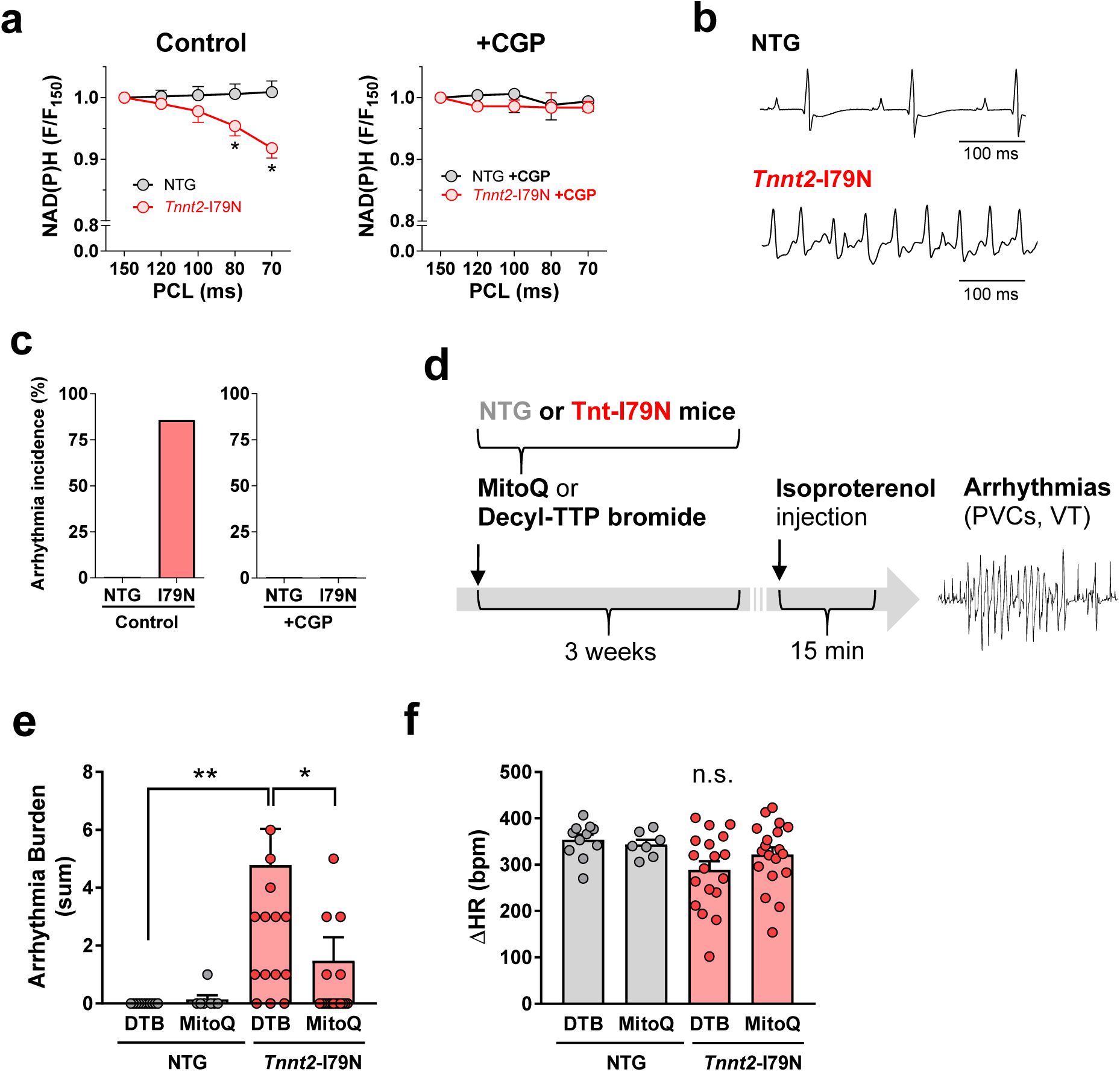
Targeting mitochondrial Ca^2+^ handling or ROS protects from cardiac arrhythmias in Tnnt2-I79N hearts *in vitro* and *in vivo* **a-c**, Hearts of *Tnnt2-*I79N or *Tnnt2*-WT mice were mounted in a Langendorff apparatus and paced at defined periodic cycle lengths (PCL) of 150 to 70 ms and NAD(P)H redox state (**a**) together with the occurrence of arrhythmias (**b, c**) were detected. Hearts were perfused with saline containing either vehicle (**a-c**; *Tnnt2*-WT, n=6; *Tnnt2*-I79N, n=7) or CGP-37157 (**c**; 10 µM; *Tnnt2*-WT, n=4; *Tnnt2*-I79N, n=5), respectively. **d**, Study protocol of NTG or *Tnnt2*-I79N mice treated with either mitochondria-targeted coenzyme Q (Mito-Q, 250 µM in drinking water; NTG, n=7; I79N, n=19) or the inactive precursor Decyl-TTP bromide (DTTP, 250 µM in drinking water; NTG, n=11; I79N, n=18), respectively. After three weeks, isoproterenol was injected in narcosis with ketamine/xylazine and PVCs and VTs recorded in the following 15 minutes. Isoproterenol-induced increase in arrhythmic burden (**f**) and heart rate (HR; **f**) in NTG or I79N mice pretreated with MitoQ or DTTP, respectively. Error bars indicate the standard error of the mean (SEM); n-numbers in the legend are indicated as numbers of hearts (**a-c**) or mice (**e-f**), respectively. Paired samples t-Test was conducted when one-matched variable was tested between the groups of data, while unpaired t-Test was performed for un-matched variables. The level of significance was set to *p<0.05.

To test whether mitochondrial therapy also works *in vivo*, we treated *Tnnt2*-I79N or NTG mice for 3 weeks with orally applied mitochondrially-targeted ubiquinone (Mito-Q, **Fig. 7d**), a well-established mitochondrial antioxidant,^28^ or its inactive precursor, Decyl-TTP bromide (DTTP). Compared to NTG, *Tnnt2*-I79N mice treated with DTTP exhibited a large burden of ventricular arrhythmias in response to Iso (1.5 mg/kg; **Fig. 7e**) despite a similar increase in heart rate (**Fig. 7f**). This is consistent with the previously reported arrhythmia phenotype of *Tnnt2*-I79N mice.^15,16^ In contrast, long-term treatment with Mito-Q substantially reduced the Iso-induced arrhythmia burden of *Tnnt2*-I79N mice (**Fig. 7e**). Taken together, these data establish proof of principle for using mitochondrial therapies to prevent ventricular arrhythmias in HCM.

## Discussion

Oxidative stress plays a central pathophysiological role in various forms of heart failure.^64^ In patients with HCM, myocardial ROS levels correlate with the severity of the disease,^65^ and reducing oxidative stress with N-acetyl cysteine (NAC) ameliorates LV hypertrophy, fibrosis and arrhythmias in preclinical models of the disease.^66,67^ Large-scale multi-omics studies on human samples from patients with advanced HCM revealed mitochondrial dysfunction and oxidative stress, including depletion and/or oxidation of NADH, NADPH and glutathione.^10–14^ However, the exact upstream mechanisms how sarcomeric variants actually cause these defects, and whether these can underlie cardiac arrhythmias, were so far unknown.

Here, we report that increased myofilament Ca^2+^ affinity, a canonical consequence of most HCM-causing mutations,^2,3^ induces energetic mismatch through an imbalance between Ca^2+^-induced Krebs cycle activation and ADP-induced acceleration of respiration in mitochondria, oxidizing NADH and FADH_2_. In response to NADH oxidation, the mitochondrial transhydrogenase (NNT) reverses and oxidizes NADPH, with the ensuing glutathione oxidation and ROS overflow providing both a *trigger* and a *substrate* for ventricular tachycardia, fibrillation and/or AV blocks, typical forms of arrhythmia in HCM,^68^ and amendable by mitochondria-targeted therapies (**Fig. 8**). Importantly, these mechanisms occur *before* any dysfunction of mitochondria *per se* evolves and therefore, represent early disease-initiating and actionable events. Although structural^69^ and functional aspects^26^ of reverse-mode NNT were addressed by previous studies, its role in HCM was so far unknown, and our findings answer the long-standing question how sarcomeric mutations cause oxidative stress and arrhythmias in HCM, opening new avenues for research and treatment options.

**Figure 8:**
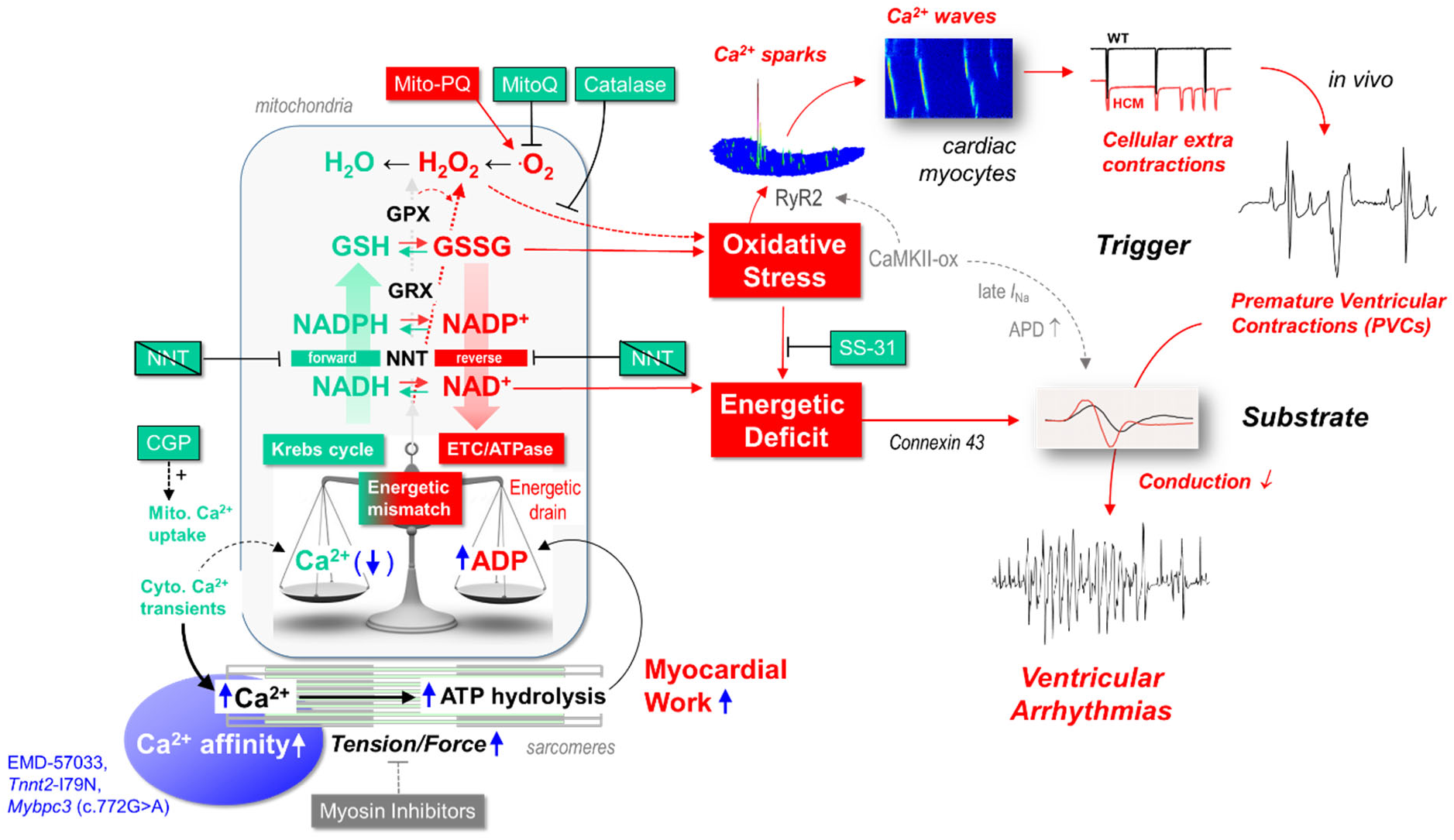
Increased myofilament Ca^2+^ affinity induces energetic and redox mismatch that serves as a trigger and substrate for arrhythmias. Increased Ca^2+^ affinity of the myofilaments, induced either by a mutation in *TNNT2* (I79N) or in *MYBPC3* (c.772G>A), or pharmacologically with EMD 57033 induce a mismatch between ADP-induced oxidation and Ca^2+^-induced reduction of the NADH and (via the mitochondrial transhydrogenase, NNT) NADPH redox state. On one hand, NADH oxidation may contribute to energetic deficit that (via dephosphorylation of connexin 43)^19^ slows atrio-and intra-ventricular electrical propagation, providing a *substrate* for arrhythmias. On the other hand, oxidative stress induced by NADPH and glutathione oxidation trigger arrhythmias by provoking spontaneous sarcoplasmic reticulum Ca^2+^ release events (“sparks”) that can result in Ca^2+^ waves, delayed depolarizations and thereby, premature ventricular contractions (PVCs) *in vivo*. Furthermore, mitochondrial ROS can aggravate the energetic deficit that accounts for slowed electrical conduction. Additional factors, not tested in this study, may be redox-dependent activation of CaMKII, which can phosphorylate RyR2 and increase late *I*_Na_ and thereby, proaction potential duration (APD). Together, these processes induce ventricular arrhythmias. Pharmacological therapies targeted at mitochondrial Ca^2+^ handling (CGP-37157), oxidative stress (MitoQ) and/or cardiolipin (SS-31) may prevent the development of arryhthmias in HCM.

### Mitochondrial ROS aggravate energetic depletion as a substrate for arrhythmias

In cardiac myocytes (**Fig. 1i**, **4b**) and whole hearts (**Fig. 7a**) of mice with HCM-typical mutations, mitochondrial NAD(P)H and FADH_2_ excessively oxidized during β-adrenergic stimulation and/or pacing. Since NADH and FADH_2_ are required for ATP production at the respiratory chain, this may reduce phosphocreatine (PCr) levels and creatinine kinase flux, as observed in patients with HCM and Ca^2+^-sensitizing mutations in *MYH7*, *MYBPC3* or *TNNT2*.^70,71^ Decreased PCr/ATP ratios in HCM reduce the calculated free energy of ATP hydrolysis (ΔG_∼ATP_), which compromises excitation-contraction coupling and the ability of the heart to respond to inotropic challenge.^72,73^ In addition, insufficient rephosphorylation of ADP aggravates diastolic dysfunction through ADP-induced elevation of myofilament force development.^74^

Furthermore, focal energy deprivation after isoproterenol injection dephosphorylates connexin 43 in *Tnnt2*-I79N mice, which slows transversal electrical conductance and thereby provides a substrate for arrhythmias *in vivo*.^19^ Here, we show that oxidative stress aggravates energy deprivation, since slowing of intra-(**Fig. 5f**) and atrio-ventricular conduction (**Fig. 5g**) in HCM hearts in response to β-adrenergic stimulation required a functional NNT. We discovered recently that in response to pathological cardiac afterload, NADH oxidation towards ATP production reverses the NNT to regenerate NADH from NADPH, depleting the anti-oxidative capacity and giving rise to H_2_O_2_ emission.^26^ When a functional NNT is missing (as in C57BL/6J mice^24^), the NADPH pool is preserved under energy-demanding conditions, preventing H_2_O_2_ emission.^26^ Conversely, pathological ATP demand caused by Ca^2+^ sensitizing mutations of the myofilaments^70^ (e.g., by *Tnnt2*-I79N^15,16,55^) provoked ROS emission and arrhythmias in cardiac myocytes *with* a functional NNT (**Fig. 4f**). In contrast, the absence of a functional NNT abrogates the cellular ROS increase and arrhythmias in *Tnnt2*-I79N myocytes (**Fig. 4f**). Since also *in vivo*, NNT deficiency ameliorated isoproterenol-induced slowing of (atrio-) ventricular conduction in *Tnnt2*-I79N hearts (**Fig. 5d-g**), these results strongly support the notion that oxidative stress, secondary to NADPH oxidation via reverse-mode NNT, aggravates the energetic deficit that promotes slowing of conduction (**Fig. 8**).

SS-31, a tetrapeptide that binds to cardiolipin in the inner mitochondrial membrane,^27^ prevented Iso-induced slowing of (atrio-) ventricular conduction only in *Tnnt2*-I79N mice *with* (but not *without*) a functional NNT (**Fig. 5d-g**), but without reducing myocardial ROS *per se* (**Fig. 5b, c**). Since SS-31 scavenges neither H_2_O_2_ (**Supplemental Fig. 5**) nor .O_2_ directly, these data suggest that *i)* SS-31 protects hearts from ROS-induced conduction delay downstream of ROS and *ii)* the previously described connexin 43-dependent focal energy deprivation in *Tnnt2*-I79N mice^19^ is at least to some extent induced by ROS-related compromise of mitochondrial ATP production (**Fig. 8**). These data are in line with the concept that SS-31 prevents oxidation of cardiolipin and thereby preserves the integrity of the ETC, avoiding electron leakage to O_2_ to form .O_2_ . Furthermore, SS-31 was reported to reduce proton leak via the adenine nucleotide transporter (ANT) and improve the efficacy of electron transfer along the ETC in aged hearts.^75^ In fact, SS-31 improved NADH-coupled respiration in *ex vivo* myectomy samples from patients with heart failure with reduced ejection fraction (HFrEF)^76^ or HCM.^14^

### Mitochondrial ROS trigger arrhythmias

In addition to providing a *substrate*, mitochondrial ROS also serve as a *trigger* for arrhythmias (**Fig. 8**). Since in the presence of the Ca^2+^ sensitizer EMD 57033, mitochondrial overexpression of catalase suppressed basal and Iso-induced ROS (**Fig. 3b**), spontaneous SR Ca^2+^ release events (“Ca^2+^ sparks”) and autonomous Ca^2+^ waves in isolated cardiac myocytes (**Fig. 3c-f**), H_2_O_2_ is the reactive oxygen species responsible for arrhythmia induction, and mitochondria its source. Furthermore, the increases in ROS (**Fig. 4f**) and arrhythmias (**Fig. 4g**) in *Tnnt2*-I79N myocytes were completely blunted in the absence of an NNT, emphasizing the central role of mitochondrial NADPH oxidation and depletion of H_2_O_2_-eliminating anti-oxidative systems^20,26,45^ as upstream mechanism. Finally, the mitochondria-targeted redox cycler Mito-PQ^58^ induced cellular arrhythmias *in vitro* (**Fig. 6i**) and premature ventricular contractions (PVCs) and tachycardia during β-adrenergic stimulation in normal (non-transgenic) hearts *in vivo* (**Fig. 6i,j**). Employing genetically encoded reporters, we could resolve that cellular arrhythmias in response to Mito-PQ were related to oxidation of the mitochondrial and cytosolic glutathione pools, which are replenished by NADPH, and the (over-) expression of glutathione reductase (Grx) coupled to these fluorescence reporters in the cytosol prevented the arrhythmias. Finally, arrhythmias in *Tnnt2*-I79N mice were abrogated by 3-week *in vivo* pretreatment with Mito-Q, a mitochondria-targeted ROS scavenger^28^ (**Fig. 7e**).

CGP-37157, which inhibits mitochondrial Ca^2+^ efflux via the NCLX and thereby fosters mitochondrial Ca^2+^ accumulation,^43,47^ prevented mitochondrial NAD(P)H oxidation, ROS emission and arrhythmias in cardiac myocytes and whole hearts of *Tnnt2*-I79N mice (**Fig. 4b-d** and **Fig. 7a-c**), in *Mybpc3*-KI myocytes (**Fig. 2a-c**) and WT myocytes pretreated with the Ca^2+^ sensitizer EM 57033 (**Fig. 2h, i**). Together, these data indicate that increased myofilament Ca^2+^ affinity causes an imbalance of mitochondrial Ca^2+^ in relation to the heightened ADP-mediated acceleration of respiration in HCM. The ensuing oxidation of NADH and FADH_2_ induces an energetic deficit, while NADPH oxidation depletes mitochondrial and cytosolic glutathione, increasing ROS that *i)* aggravate the energetic deficit and *ii)* trigger arrhythmias. These processes conspire to trigger and sustain arrhythmias *in vivo* (**Fig. 8**).

### How mitochondrial ROS trigger and maintain arrhythmias

A large body of evidence supports the notion that pathologically elevated ROS cause arrhythmias.^51^ High amounts of ROS increase diastolic [Ca^2+^]_c_ by oxidizing RyR2^52,53^ and inhibiting SR Ca^2+^-ATPase.^77–79^ This activates the sarcolemmal Na^+^/Ca^2+^ exchanger and Ca^2+^- induced Ca^2+^ release from the SR, eliciting abnormal action potentials that trigger arrhythmias.^80^ In fact, close proximity of mitochondria to the SR facilitates this regulatory crosstalk of ROS and Ca^2+^ in cardiac myocytes.^36,80^ In particular, oxidation of cytosolic glutathione, as we observed in response to Mito-PQ, oxidizes RyR2 and triggers spontaneous Ca^2+^ release events,^81,82^ similar to the ones we observed in response to the Ca^2+^ sensitizer EMD 57033 and which were blunted by mitochondrial catalase overexpression.

These proarrhythmic effects may be further aggravated by ROS-induced activation of Ca^2+^/calmodulin-dependent protein kinase II (CaMKII),^83,84^ which phosphorylates Na^+^ channels, elevating the late Na^+^ current (*I*_Na,L_), increasing intracellular [Na^+^] and prolonging action potential duration.^85^ Together with (ROS- and) CaMKII-induced activation of L-type Ca^2+^ channel currents (*I*_Ca,L_), this can additionally cause early (EADs) and delayed afterdepolarizations (DADs).^86^ In fact, in cardiac myocytes isolated from patients with HCM (with various gene defects), many of these alterations of EC coupling could be reconciled, such as elevated CaMKII activity, enhanced *I*_Na,L_ and *I*_Ca,L_, prolonged action potential duration and increased occurrence of cellular arrhythmias (**Fig. 8**).^87^

### Mitochondrial ROS could also underlie maladaptive cardiac remodeling in HCM

Besides the induction of arrhythmias, ROS and glutathione oxidation may also contribute to maladaptive remodeling, since LV hypertrophy^66^ and fibrosis^66,67^ in animal models of HCM could be reversed by the nonspecific antioxidant NAC, which reduces glutathione. The force-time integral, which is increased in HCM, but reduced in hereditary forms of *dilated* cardiomyopathy, predicts the development of concentric LV hypertrophy.^40^ In this model, concentric LV hypertrophy (but not the mere increase in myocardial mass) is driven by the mitogen-activated protein (MAP) kinase ERK1/2,^40^ which is activated in HCM in a ROS-dependent manner.^88^ Therefore, it may be speculated that the newly identified mechanism of increased mitochondrial ROS emission also determines the mode of LV hypertrophy (eccentric versus concentric^40^) in hereditary forms of cardiomyopathy in the long term. In this context, it is noteworthy that genetic or pharmacological targeting of mitochondrial, but not cytosolic ROS reduced hypertrophy and fibrosis, also typical hallmarks of remodeling in HCM, in a murine diastolic heart failure model.^31,89^ Therefore, future studies should address the therapeutic potential of mitochondrially-targeted drugs on cardiac remodeling in HCM.

### Clinical implications for the treatment of HCM

The main clinical manifestations of HCM are dyspnea, angina and SCD caused by ventricular arrhythmias. While β-blockers, the Ca^2+^ channel blockers diltiazem and verapamil as well as cardiac myosin inhibitors improve symptoms in patients with HCM,^4,90–92^ no medical treatment efficiently reduces the risk of SCD, limiting its efficient prevention to implanted cardiac defibrillators.^4,93,94^ Here, we show that in two different mouse models of HCM, four different interventions that target mitochondria (NCLX inhibition with CGP, genetic NNT inactivation, cardiolipin protection with SS-31 and ROS scavenging with MitoQ) prevent arrhythmias in cardiac myocytes, isolated hearts and *in vivo* (**Fig. 8**).

In preclinical studies, NCLX inhibition with CGP^95^ and mitochondrial ROS scavenging with Mito-TEMPOL^96^ reduced arrhythmias and maladaptive remodeling in a guinea pig model of HFrEF. While CGP was never tested clinically, the Ca^2+^ channel blockers verapamil and diltiazem both have NCLX-inhibiting properties.^97^ MitoQ consists of ubiquinone (also known as coenzyme Q), an electron carrier in the mitochondrial respiratory chain,^98^ coupled to the lipophilic cation TPP^+^.^28^ Coenzyme Q itself was tested in numerous smaller clinical trials on patients with HF.^99^ In the largest of these, coenzyme Q reduced cardiovascular and total mortality in 420 patients with HFrEF.^100^ In a smaller, open-label study on patients with HCM, coenzyme Q improved cardiac hemodynamics and symptoms.^101^ However, there is still little clinical data for the *conjugated* MitoQ, which is safe in humans and, similar to coenzyme Q, sold over the counter on the free market. One ongoing study investigates the effect of MitoQ on exercise capacity in patients with HF with preserved ejection fraction (HFpEF) and chronic kidney disease (NCT03960073).

SS-31, also known as Elamipretide, has already been tested in humans with various diseases. Although in patients with HFrEF, a single intravenous dose modestly improved cardiac volumes,^102^ no significant effects on cardiac hemodynamics were noted after a 4-week subcutaneous treatment.^103^ Barth syndrome is a rare mitochondrial cardiomyopathy which typically manifests as dilated cardiomyopathy in the first months of life and then transitions into a phenotype of HFpEF or even HCM.^44,104,105^ In these patients, elamipretide reversed cardiac remodeling and improved LV stroke volume after 3-year open label treatment compared to baseline, and improved the 6-minute walk distance after 36 and 168, but not 12 weeks of treatment.^105,106^

Based on our observations that the absence of a functional NNT prevented oxidative stress and arrhythmias, this protein may represent another attractive pharmacological target. NNT protein is robustly expressed in human heart and modestly downregulated in patients with HFrEF.^107^ Although mutations in the *NNT* gene are associated with adrenocortical deficiency^108,109^ and left ventricular non-compaction cardiomyopathy,^110^ C57BL/6J mice, which are lacking a functional NNT, have a normal heart function and lifespan, but are protected from HF induced by left^26^ or right ventricular pressure overload^111^ or a combination of high fat diet and the nitric oxide synthase inhibitor L-NAME,^112^ the currently most commonly used HFpEF model.^113^ Cryo-electron microscopy-based resolution of the structure of intact mammalian NNT in different conformational states, including the reverse mode, may facilitate the development of small compounds directed against the NNT.^69^

Cardiac myosin inhibitors reverse the pathologically increased actin-myosin interaction in HCM^114^ and improve hemodynamics and symptoms,^90,92^ improve quality of life^115^ and reverse cardiac remodeling^116,117^ in patients with obstructive, but also non-obstructive HCM.^91, 118^ According to the concept presented here, myosin inhibitors should also reduce energetic mismatch, oxidative stress and arrhythmias from upstream (**Fig. 8**).

Together, our work may therefore serve as a mechanistic framework for the development of novel therapeutic regimens in patients with HCM by facilitating the synergistic profiles and clinical safety of the mentioned drugs (**Fig. 8**).

## Acknowledgements

We thank Michelle Gulentz and Nina Schnellbach for technical assistance, Peter S. Rabinovitch (University of Washington, Seattle, WA, US) for providing mCAT mice, and Rüdiger Pryss (Institute for Clinical Epidemiology and Biometrics, Würzburg University) for statistical consultancy. Support was and is provided by the Deutsche Forschungsgemeinschaft (DFG) to C. M. (Heisenberg Professur; SFB 894, TRR-219 and SFB 1525 (project # 453989101); Ma 2528/7-1 and 8-1 (project #505805397)), D.M.K. (RTG 2824), M.H., L.P.R. and M. B. by Project-ID 322900939 (SFB TRR-219), and A.D. by DI 2876/2-1 (#519332007). C.M. was supported by the Corona Stiftung, Margret Elisabeth Strauß-Projektförderung of the Deutsche Herzstiftung and the Federal Ministry of Education and Research (BMBF; 01EO1504, CF.3 and RC.2). L.C. and T.E. are supported by the DZHK (German Centre for Cardiovascular Research) and the German Ministry of Research Education (BMBF). F.W.F and F.F. received support from Deutsche Stiftung für Herzforschung (F/28/12, F/24/15 and F/15/17), and F.F. is supported by the DFG, project 316865582). B.C.K. is supported by U.S. National Institute of Health Grants HL071670, HL124935, and HL128044.

## AUTHOR CONTRIBUTIONS

C.M., B.C.K., T.E., and L.C. designed the study and wrote the manuscript. M.K., S.P., A.D., V.S., O.R., J.B., Q.T., F.W.F, F.F., M.S.N, J.S., D.K., A.G.N., F.A., R.V., F.J.B., R.K., V.S., E.B., L.P.R., A.K. and M.H. performed the experiments, data analysis and gave valuable scientific input. U.L., D.K., M.H., M.B. and P.L. gave valuable scientific input and contributed to editing the manuscript.

