## Supplemental Material for "Mitochondrial reactive oxygen species cause arrhythmias in hypertrophic cardiomyopathy"

### Supplementary Methods

#### *In vitro* experiments

##### Isolated cardiac myocytes

Cardiac myocytes were isolated by enzymatic digestion and experiments performed as described previously.[1,2] Briefly, adult ventricular myocytes were electrically stimulated at 0.5 Hz in Normal Tyrode´s solution containing (in mM) NaCl 130, KCl 5, $MgCl_2$ 1, Na-HEPES 10, $CaCl_2$ 1, glucose 10, pH 7.4 and then exposed to isoproterenol (30 nM) and an increase of stimulation rate to 5 Hz for 3 min. These conditions apply to experiments on *Tnnt2*-WT and -I79N mice (**Fig. 4**) and on C57BL/6N mice in the presence of CGP-37157 (10 µmol/L; **Fig. 2d-i**). In experiments on myocytes from *Mybpc3*-KI and corresponding WT mice, a shorter stimulation protocol (1 min) and lower isoproterenol concentration (10 nM) were chosen to avoid premature death of cardiomyocytes, since *Mybpc3*-KI myocytes were particularly sensitive to the stress conditions and would die if the stimulation protocol exceeded one minute (**Supplemental Fig. 2g** and see also our previous data[3]). For the experiments on mCAT mice, myocytes were stimulated at 0.5 Hz for 1 min, then EMD 57033 (3 µM) was washed in for 4 min and then isoproterenol (30 nM) for 1 min, before stimulation rate was increased to 5 Hz for 3 min (**Fig. 3a, b**).

Sarcomere shortening was detected together with either *i)* the redox state of $NAD(P)H/NAD(P)^+$ and $FADH_2/FAD$ by autofluorescence, *ii)* $\Delta\Psi_m$ (with tetramethylrhodamine methyl ester; TMRM) together with $[Ca^{2+}]_c$ (indo-1 acetoxymethyl esther; AM), *iii)* mitochondrial $\cdot O_2^-$ (MitoSOX) or *iv)* cellular ROS formation (using 5-(-6)-chloromethyl-2,7-dichlorohydrofluorescein di-acetate; CM-$H_2$DCFDA), as described previously.[1,2,4,5] Calibration of $NAD(P)H/NAD(P)^+$ and $FADH_2/FAD$ was performed by applying FCCP (5 µM) and cyanide (4 mM) at the end of every experiment. To detect mitochondrial $\cdot O_2^-$, myocytes were loaded with MitoSOX (3.3 µmol/L) for 30 minutes at 37°C. The excitation of MitoSOX at 380 nm (with emission at 580 nm) allows a rather specific detection of 2-hydroxy-ethidium over ethidium, where 2-hydroxy-ethidium is a specific product of oxidation by $\cdot O_2^-$, while ethidium is not selective for oxidation by ($\cdot O_2^-$), but also sensitive to other ROS.[6,7] As a positive control, antimycin A (150 µmol/L) was used. Calibration of CM-$H_2$DCFDA was performed by applying 10 mM of extracellular $H_2O_2$. Only myocytes with a clear increase in CM-$H_2$DCFDA fluorescence in response to $H_2O_2$ were used for analysis, and no differences were observed in baseline DCF fluorescence or the response to extracellular $H_2O_2$ between groups. Using a patch-clamp based protocol with application of indo-1 salt and rhod-2 AM, $[Ca^{2+}]_c$ was determined together with $[Ca^{2+}]_m$ in *Mybpc3*-KI and WT myocytes under similar conditions.[1,2,5,8]

For the analysis of cellular arrhythmias, the number of unstimulated beats (systolic sarcomere shortenings) were quantified within the first 20 s after stepping stimulation frequency from 5 Hz to 0.5 Hz (i.e., 10 regular beats per cell) in myocytes that were not loaded with any dye (i.e., the cells that were used for the analysis of $NAD(P)H/NAD(P)^+$ and $FADH_2/FAD$ autofluorescence). During this time, the solution was switched back to normal Tyrode´s solution without isoproterenol, but the bath was not completely cleared of isoproterenol at this time yet.

In cardiac myocytes of mice expressing mito-roGFP2-Orp1, cyto- or mito-Grx1-roGFP2, excitation was alternated between 340 nm and 405/490 nm, and emission collected at 450 nm and 520 nm to determine mitochondrial NAD(P)H and $H_2O_2$ (mito-roGFP2-Orp1) or oxidized glutathione (GSSG; cyto- or mito-Grx1-roGFP2) levels within the same cells, respectively. Mitochondria-targeted paraquat (Mito-PQ) was applied at 10 µM.

### Confocal microscopy for $Ca^{2+}$ sparks and $Ca^{2+}$ waves analyses

Cardiac myocytes were loaded with 1 µM Fluo-8/AM (1 µl of a stock in which Fluo-8/AM was solved in dimethyl sulfoxide, DMSO, complimented with 20% Pluronic F-127) in Tyrode´s solution for 30 min at room temperature, followed by 15 min in dye-free Tyrode´s solution for de-esterification. Cells were pretreated with EMD 57033 (EMD; 3 µM) or a similar amount of DMSO (control) for 4 min and then mounted in a custom-made chamber onto the stage of a motorized Nikon microscope Eclipse Ti (NIKON, Tokyo, Japan), perfused with Tyrode´s solution at 35°C. EMD or vehicle were present for the remainder of the experiment, respectively.

A high-speed 2D-array confocal scanning module, VT-Infinity3 (Visitech International, Sunderland, United Kingdom), was equipped with a Hamamatsu ORCA-Flash 4.0 sCMOS camera (Hamamatsu Photonics, Hamamatsu City, Japan) and coupled to the microscope. Solid-state laser lines at 488 nm (Cobolt AB, Solna, Sweden) were guided into the confocal module via a light guide. AOTF of the Laser was set to 25%. The scanning speed of the confocal module was set to 600 Hz, 60% of the chip area of the camera was used at its highest readout speed resulting in a recording speed of 147 images/second. The cells were imaged with Nikon Plan Fluor 60X oil DIC objective (NA=1.40, NIKON, Tokyo, Japan). The built-in dichroic mirror in the confocal scanning head VT-Infinity was a ZT488/640rpc (Chroma Technology Corporation, Vermont, USA). The emission fluorescence of Fluo-8 from 500 nm to 620 nm was collected. The camera operated in its 2x2 binning mode yielding a pixel size of 215 nm. More detailed instrument configurations and data processing procedures were described previously.[9] To discriminate $Ca^{2+}$ waves from sparks, a specific threshold of the $90^{th}$ percentile of the FWHD of all signals (=9.1 µm) was applied.

The cells were first stimulated in an electrical field (MyoPacer; IonOptics, Westwood, MA, USA) for 1 min at 0.5 Hz and then imaged for 6.8 s without any stimulation. Thereafter, myocytes were stimulated at 0.5 Hz and exposed to 30 nM isoproterenol for 30 s and then recorded without stimulation for 6.8 s. Finally, myocytes were paced at 5 Hz for 60 s and again, imaged for 6.8 s.

### Isolated mitochondria

Mitochondria were isolated from *Mybpc3*-KI and WT mouse hearts as described previously.[2] Mitochondria were supplied with pyruvate/malate as substrates (5 mM each). Respiration (Clark electrode), $\cdot O_2^-$ formation (electron paramagnetic resonance, EPR), and $H_2O_2$ emission (Amplex Ultra-Red) were determined in the absence or presence of ADP (1 mM) as described previously.[2]

### PCR analysis for genotyping for wild-type and truncated *Nnt*

Mouse genomic DNA (gDNA) was prepared from whole tissue samples (ear or tail) using the DNA Extraction Solution 1.0 (Biozym, Cat No.101094 or QE09050) following the manufacturer´s protocol. Two µl of gDNA were inserted into PCR. While C57BL/6N mice express a full length *Nnt* gene, C57BL/6J mice have two truncated *Nnt* alleles, in which exons 7-11 are missing.[2,10] For genotyping full-length (wild-type, wt) and truncated (t) Nnt alleles, we performed two different PCR reactions in one assay. The first used primers that target a sequence occurring in exon 8 of the *Nnt* gene ("in"). This product of 157 bp is only amplified in full-length *Nnt* alleles, but not in truncated Nnt alleles stemming from C57BL/6J. The second PCR reaction used primers that span exons 6-12 of genomic DNA ("out"). In truncated *Nnt* alleles, this results in a 546 bp product, while in full-length *Nnt* alleles, the product would be far too large to become amplified (due to very long introns between exons 6 and 12 that are missing in the truncated allele). Therefore, the 546-bp product occurs only in truncated *Nnt* alleles, while no amplification product is gained in full-length WT alleles. gDNA of C57BL/6J and C57BL/6N mice were used as (homozygous) positive and negative controls for truncated and full-length *Nnt* alleles, respectively.

The primer sequences used to detect the wild-type *Nnt* gene ("in") were as follows:

- NNT_exon_8_forward: 5'-TATTGGCTACACAGACCTTCC-3' and
- NNT_exon_8_reverse: 5'-TGACGTGACTCATTGTACCA-3'.

The primer sequences used to detect the 546-bp product in the truncated *Nnt* gene ("out") were as follows:

- Nnt_exon_6-12_forward: 5'-GTAGGGCCAACTGTTTCTGC-3'
- Nnt_exon_6-12_reverse: 5'-TCCCCTCCCTTCCATTTAGT-3'

Amplification was carried out using a Primus Advanced 96 PCR Cycler (Peqlab, Erlangen, Germany) and the Taq Polymerase Kit (Peqlab; Order No. 01-1030) inserting 20 pmol/µL of each primer pair. The PCR cycling were as follows: Step 1: 95 °C for 5 min followed by 35 cycles of denaturation at 95 °C for 1 min, annealing at 60 °C for 1 min and amplification at 72 °C for 1 min. PCR was terminated by amplification at 72 °C for 10 min and PCR-products were stored at 4 °C until their separation on a 2% agarose electrophorese gel.

### Langendorff perfusion experiments

*Tnnt2*-I79N and -WT mice were deeply anesthetized in a recumbent position with 5% isofluorane in oxygen ($O_2$). After thoracotomy, the heart was rapidly excised and immersed in cold Tyrode's solution containing heparin. The aorta was cannulated with a custom-made plastic cannula and knotted with silk suture to facilitate retrograde perfusion in Langendorff mode. The cannula was then connected to a constant pressure-perfusion system with Tyrode's solution warmed to 37 ºC and bubbled with 95% oxygen and 5% carbon dioxide. The Tyrode's solution contained 130 mM NaCl, 4 mM KCl, 23 mM $NaHCO_3$, 1.5 mM $NaH_2PO_4$, 1 mM $MgCl_2$, 2 mM $CaCl_2$, 10 mM glucose and 10 µM propanolol (either Sigma Aldrich, MO, US; or Fisher Scientific, PA, US).

A platinum electrode was placed at the apex to pace the heart at 3 times threshold stimulation current. Hearts were subjected to a pacing protocol that previously has been shown to consistently trigger ventricular tachycardia in explanted *Tnnt2*-I79N hearts.[11] Briefly, starting with a pacing cycle length (PCL) of 150 ms (reflecting a heart rate of 400/min), the cycle length was shortened to 120, 100 and 80 ms (=750/min), respectively, until sustained VT was induced. In cases in which 1:1 capture (1 heart beat for 1 electrical pacing pulse) was lost prior to arrhythmia induction, the current was slowly increased until capture resumed and the increased current was recorded. For hearts that did not encounter arrhythmias by 80 ms, the PCL was progressively further stepped down in 10 ms decrements to 60 ms until arrhythmias were induced or capture was lost.

### Optical imaging of NADH in isolated hearts

All imaging work was performed on a previously-described whole-heart imaging system.[11] The system consists of a fully jacketed constant pressure Langendorff-perfusion system interfaced with a high-speed CCD camera and different excitation light sources. The Langendorff-perfused hearts were allowed to stabilize during 15 minutes of perfusion and then subjected to the pacing protocol described above. Following one minute of pacing at each PCL, hearts were briefly illuminated at 395 nm. The resulting NADH autofluorescence was optically filtered through a 450 nm filter and recorded on a 1000 frame per second, 80x80 pixel CCD camera (Redshirt, GA, US). Data were analyzed offline in MATLAB (Mathworks, MA, US) by selecting a 10x10 pixel window over a well-perfused region of the left ventricle and averaging the fluorescence signal. As calibrating the signal was not possible, data are presented as $F/F_0$. In CGP treated hearts, 10 µM of CGP was recirculated for 10 minutes prior to the start of pacing.

### *In vivo* experiments

Studies were approved by the Vanderbilt Institutional Animal Care and Use Committee. Age-matched male and female *Tnnt2*-I79N and NTG mice were used for all studies. *Tnnt2*-I79N mice heterozygous for truncated nicotinamide nucleotide transhydrogenase (*Nnt*) allele were maintained on an outbred mouse strain B6SJLF1/J (100012). This breeding strategy resulted in the generation of *Tnnt2*-I79N and NTG mice with variable *Nnt* status including wt/wt, wt/t or t/t, with wt/wt denoting two wild-type alleles and t/t denoting the presence of two truncated alleles and therefore, the absence of a functional NNT. Age and biological replicates are as indicated for the respective experiment.

### Electrocardiogram (ECG) in anesthetized mice

For the experiments in **Fig. 5a**, a total of 68 age-matched mice (9-12 week-old; n=37 NTG and n=31 *Tnnt2*-I79N) of both genders were used for experiments. Mice were anesthetized with inhaled isoflurane (4% for induction, 2% for maintenance) in 100% $O_2$ with a flow of 2 L/min, while spontaneously breathing and placed in prone position on a warm surface as described previously.[12] Surface ECG was obtained by placement of subcutaneous 24-gauge electrodes into all four limbs and connected to ECG leads. ECG (Leads I and II) were recorded using animal bioamplifiers linked to a PowerLab station (AD Instruments) and LabChart5 software. The surface ECG was monitored continuously and stored on disk. P, Q, R and S waves as well as PR and QRS intervals were identified as shown in a representative example in **Supplemental Figure 7a**. ECG intervals (RR, PR, QRS; seconds) and amplitudes (RS; millivolts) were measured in standard fashion.

### SS-31 treatment and ECG protocol

For the experiments in **Fig. 5**, one hour prior to ECG recordings, NTG (n=37) and *Tnnt2*-I79N animals (n=31) were randomly assigned to either receive an intraperitoneal (i.p.) injection of vehicle solution (0.9% NaCl; NTG: n=20; I79N: n=17) or SS-31 (3 mg/kg; NTG: n=17; I79N: n=14), respectively. After a baseline ECG was obtained (~2 min), isoproterenol (Iso, 3 mg/kg) was injected i.p. and the ECG recordings continued for at least 8 min. The effect of isoproterenol was recorded at 1, 4 and 8 min after the iso injection (**Fig. 5a**). In **Fig. 5d-i**, the analyses are related to the respective *Nnt* genotypes (wt/wt or wt/t versus t/t; for PCR protocols, see above).

### Electrocardiogram analysis

Electrocardiogram (ECG) recordings were obtained using needle electrodes connected to PowerLab (ADInstruments) for downstream analysis using Lab Chart software. Body temperature was maintained using microwavable gel warming pad monitored by surface thermometer. PR, QRS, and heart rate were manually assessed using the mark-measure feature on LabChart. ECG records were analyzed blinded for genotype and drug treatment. RR, PR and QRS intervals, and RS amplitude were determined by averaging 5 consecutive beats during sinus rhythm during baseline and 1, 4 and 8 min after isoproterenol administration, respectively (**Supplemental Fig. 7a, b**). Since mice do not show a clear flat ST-segment as humans, the S-peak was used to determine QRS duration. Recordings were also examined for the presence of premature ventricular contractions (PVCs), couplets (2 consecutive PVCs) and ventricular tachycardia (3 or more consecutive PVCs).

To quantify ventricular arrhythmia burden for each mouse, we used a scoring system adapted from Curtis and Walker (**Supplemental Table 1**).[13] The summative score (sum of scores reached for individual mouse) was used as primary outcome measure of ventricular arrhythmia burden.

### Immunofluorescence analysis of 8-Hydroxyguanosine

For the analysis of oxidative stress in the myocardium of *Tnnt2*-I79N and NTG mice displayed in **Fig. 5B**, from a subset of the hearts displayed in **Fig. 5a** (Vehicle: NTG, n=9; I79N, n=7; SS-31: NTG, n=7; I79N, n=7), deparaffinised tissue sections (3 µm thick) were placed in Coplin jars with 0.05% citraconic anhydride solution, pH 7.4, for 1 hour at +98°C and then incubated overnight at 4°C and for 3 hours at 37°C the next day with the first antibody, followed by the appropriate secondary antibody at 37°C for 20 minutes. Immunofluorescence studies were performed by applying a polyclonal antibody against 8-Hydroxyguanosine (abcam, ab10802, UK), diluted 1:300 with the mixture of citrate buffer (4xSSC) and 0.1% Tween 20 to detect oxidative stress in the cells. Monoclonal antibody against α–sarcomeric actin (clone5c5, Sigma-Aldrich, Germany) was used to detect cardiomyocytes. FITC- and TRITC-conjugated anti-goat IgG and anti-mouse IgM, (all Dianova, Germany) were used as secondary antibodies. To perform wash steps, 4xSSC buffer was used. Sections were counterstained with DAPI (Calbiochem, Germany) and mounted with fluorescent mounting medium (Vectashield, Vector Laboratories, Burlingame, CA, USA) for fluorescence microscopic analysis. All sections were analyzed using a Nikon Eclipse Ni epifluorescence microscope (Nikon, Germany) with appropriate filters.

Numbers of 8-hydroxyguanosine-positive cardiomyocytes per $mm^2$ were determined by examining sections double stained for 8-Hydroxyguanosine and α-sarcomeric actin, respectively, in 15 randomly chosen fields at 1000x magnification, (1 section per heart). Nuclei positive for α-sarcomeric actin and double positive for 8-hydroxyguanosine and α-sarcomeric actin were counted. The cell density was calculated using the formula: cell density ($mm^2$)=cell number in 15 fields/(0.00153664)*15, where 0.00153664 is the area of the sampling grid at 1000x magnification, and 15 is the number of counted fields.

### Pharmacological induction of mitochondrial ROS with Mito-Paraquat

10 mM stocks were generated with 5 mg Mito-PQ using 523.84 µl DMSO and stored at -20°C. Prior to use, a new aliquot was diluted with 1 mL of 0.9% saline for a final concentration of 100 µM. Mice were injected with 25 µl of this solution for delivery of 2.5 nmol MitoPQ or up to 100 µl for 10 nmol.[14]

Twelve NTG mice aged 10-20 weeks were administered either 2.5 nmol Mito-Paraquat (Mito-PQ; n=5; Abcam: ab146819) or vehicle (0.9% saline; n=7) 20 minutes prior to anesthesia with ketamine (100mg/kg) and xylazine (2.5mg/kg) IP. After 2 minutes of baseline ECG recording, mice received 1.5 mg/kg isoproterenol (in 0.9% saline) IP followed by up to 30 minutes of additional ECG monitoring. High dose MitoPQ studies involved baseline ECG recording for 2 minutes followed by administration of either vehicle or 10 nmol MitoPQ with up to 30 minutes of additional ECG monitoring.

### Evaluation of anti-arrhythmic effects of the mitochondrial ROS scavenger MitoQ

Non-transgenic (NTG; n=18) and *Tnnt2*-I79N mice (I79N; n=37) at an age of 8-13 weeks were administered MitoQ (250 µmol/L; NTG, n=7; I79N, n=19) or the inactive precursor Decyl-TTP bromide (DTB, 250 µmol/L; NTG, n=11; I79N, n=18) in normal drinking water for 3 weeks prior to isoproterenol pharmacological challenge.[15-17] Stock solutions of both MitoQ (MedKoo Biosciences #317102) and Decyl-TTP bromide (MedKoo Biosciences #620110) were generated by dissolving compounds in 1:1 ETOH/$H_2O$. Additional compound was provided by William Stow from MitoQ Ltd (New Zealand). Samples were gently vortexed at room temperature and frozen at -20°C for use within 3 weeks of preparation. Drinking water was replenished every 3 days by fresh preparation of 250 µM MitoQ or DTB solutions using frozen stocks and autoclaved water.

Mice were anesthetized with ketamine (100 mg/kg) and xylazine (2.5mg/kg) IP. After 2 minutes of baseline ECG recordings, mice received 1.5mg/kg isoproterenol (in 0.9% saline) IP followed by up to 15 minutes of additional ECG monitoring. At death or the end of the protocol, the heart was removed, rinsed in room temperature zero calcium Tyrode's solution during separation of atrium and ventricle, and immediately frozen in liquid nitrogen.

### Chemical agents

SS-31 was obtained from GenScript, USA. EMD 57033 was provided by Merck, Darmstadt, Germany. CGP-37157 was obtained from Sigma Aldrich (C8874), Munich, Germany. All other agents were obtained from Sigma Aldrich, Munich, Germany, unless indicated otherwise.

### Supplementary Figures

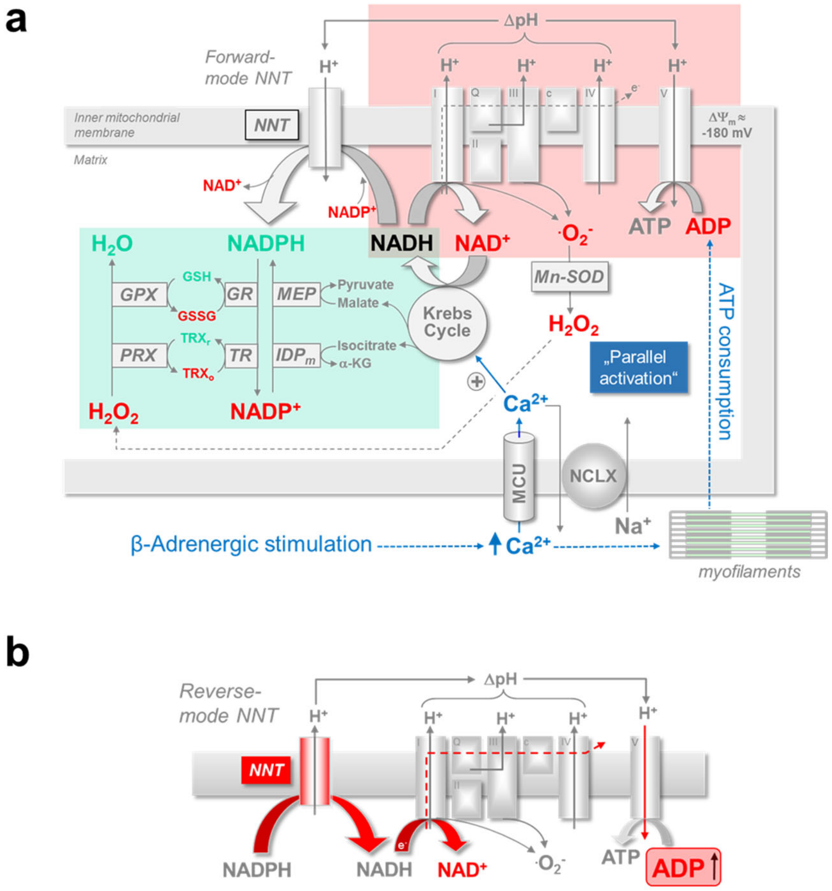


**Supplemental Figure 1:** Energy supply-and-demand matching in normal hearts

**a**, Complimentary regulation of the mitochondrial redox state by $Ca^{2+}$ and ADP. During β-adrenergic stimulation, increased cytosolic $Ca^{2+}$ transients and contraction increase ATP consumption at the myofilaments. The ensuing elevation in ADP then accelerates respiration and consumes NADH. At the same time, $Ca^{2+}$ enters mitochondria and activates Krebs cycle dehydrogenases, accelerating regeneration of oxidized $NAD^+$ to NADH. This “parallel activation” of energy consuming- and -regenerating processes maintains a reduced NADH/$NAD^+$ redox state and constant ATP/ADP ratios during varying workloads. Krebs cycle products regenerate NADPH via the nicotinamide nucleotide transhydrogenase (NNT), malic enzyme (MEP) and isocitrate dehydrogenase ($IDP_m$). NADPH then regenerates antioxidative enzymes glutathione reductase (GR) and thioredoxin reductase (TR), which in turn reduce glutathione peroxidase (GPX) and peroxiredoxin (PRX), respectively. These eliminate hydrogen peroxide ($H_2O_2$) to water ($H_2O$). **b**, When ADP-induced acceleration of respiration outweighs NADH regeneration by the Krebs cycle, NADH/$NAD^+$ becomes oxidized. We observed previously that with strong NADH oxidation (such as during high cardiac afterload), the NNT reaction can reverse, oxidizing NADPH for NADH and ATP production, but at the cost of the anti-oxidative capacity.[2]

$\Delta\Psi_m$, mitochondrial membrane potential; Mn-SOD, $Mn^{2+}$-dependent superoxide dismutase; $TRX_{r/o}$, reduced/oxidized thioredoxin; GSH/GSSG, reduced/oxidized glutathione; α-KG, α-ketoglutarate; MCU, mitochondrial $Ca^{2+}$ uniporter; NCLX, mitochondrial $Na^+/Ca^{2+}$ exchanger.

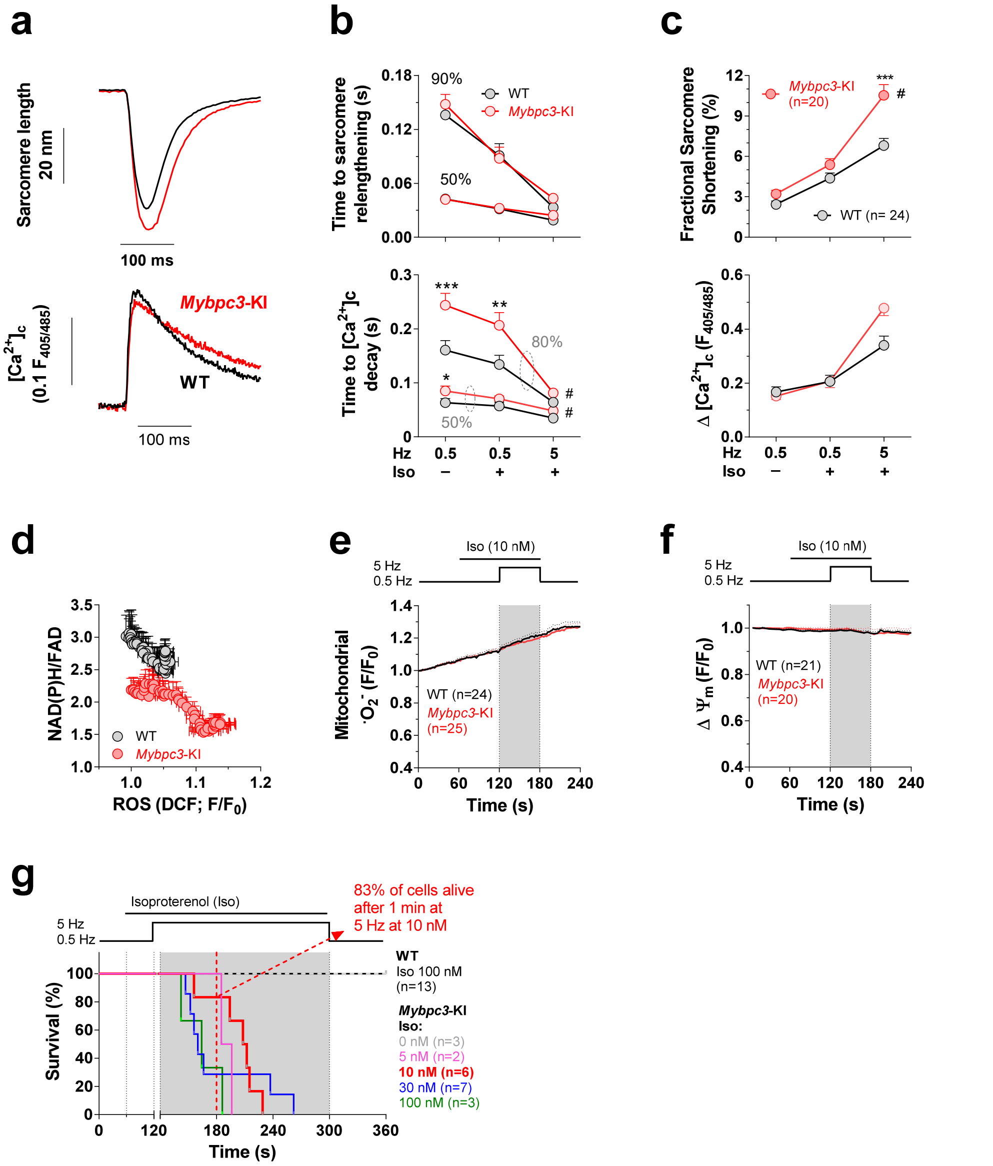


### **Supplemental Figure 2:** Excitation-contraction coupling and mitochondrial redox regulation in *Mybpc3*-KI and WT myocytes.

Experiments were performed in wild-type (WT) and *Mybpc3*-KI myocytes and the following data were obtained: **a**, Averaged original sarcomere shortening (*upper trace*; WT, n=24; *Mybpc3*-KI, n=20) and cytosolic $Ca^{2+}$ transient traces (*lower trace*; WT, n=22; *Mybpc3*-KI, n=18). **b**, Quantification of time to 50% and 90% sarcomere relengthening (*upper traces*) and time to 50% and 80% $[Ca^{2+}]_c$ decay (*lower traces*), respectively. **c**, Systolic fractional sarcomere shortening (*upper trace*) and amplitude of the systolic $[Ca^{2+}]_c$ transient (*lower trace*). **d**, NAD(P)H/FAD ratios (WT, n=25/2; *Mybpc3*-KI, n=20/2) plotted against ROS emission (indexed by DCF fluorescence; WT, n=43/6; *Mybpc3*-KI, n=24/5). **e**, Mitochondrial superoxide ($\cdot O_2^-$) formation, determined with MitoSOX. **f**, Mitochondrial membrane potential ($\Delta\Psi_m$; indexed by TMRM). **g**, Survival of WT and *Mybpc3*-KI myocytes in the presence of 0-100 nM isoproterenol, respectively.

Error bars indicate standard errors of the mean (SEM); n-numbers are indicated as numbers of cells. $^{\#}p<0.05$ (2-way ANOVA), $^{*}p<0.05$, $^{**}p<0.01$, $^{***}p<0.001$ (Bonferroni post-test).

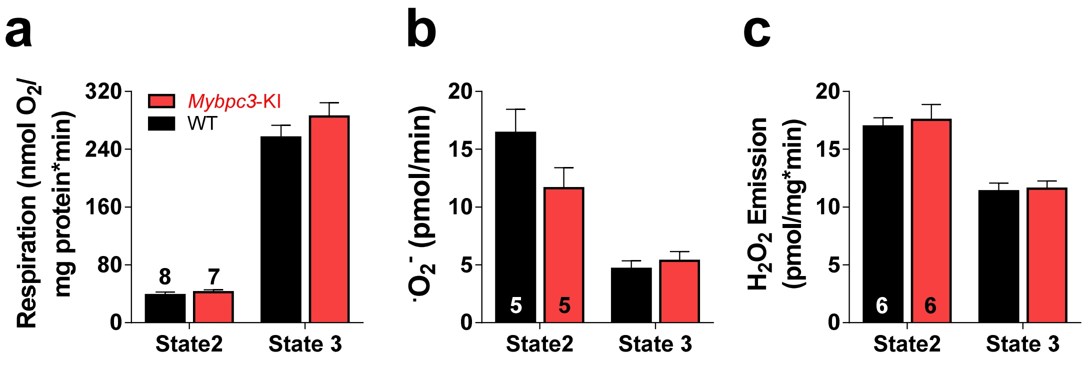


**Supplemental Figure 3:** Mitochondrial respiration and ROS production in isolated mitochondria of *Mybpc3*-KI and WT hearts

Experiments were performed on mitochondria isolated from hearts of WT or *Mybpc3*-KI mice and respiration (**a**), superoxide formation ($\cdot O_2^-$, **b**) and $H_2O_2$ emission (**c**) were determined in the absence (State 2) or presence of 1 mM ADP (state 3), respectively.

Error bars indicate standard errors of the mean (SEM); n-numbers are indicated as numbers of hearts.

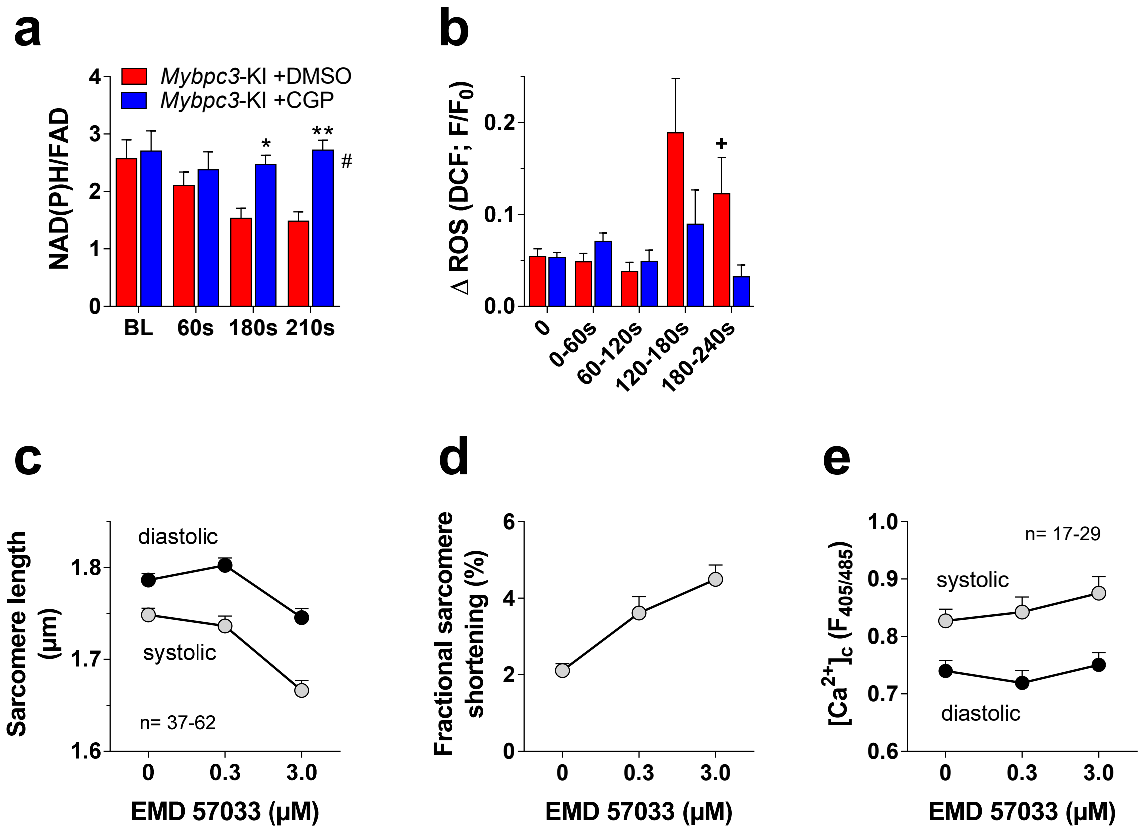


**Supplemental Figure 4:** Mitochondrial redox state and ROS emission in *Mybpc3*-KI myocytes, and the effects of the $Ca^{2+}$ sensitizer EMD 57033 on sarcomere shortening and $[Ca^{2+}]_c$ in normal C57BL/6N myocytes

**a** and **b**, Experiments were performed on *Mybpc3*-KI myocytes and the ratio of NAD(P)H/FAD was quantified at baseline (BL), 60s, 180s and 210s in the presence of either CGP-37157 (CGP, 10 µM; n=11) or vehicle (DMSO; n=11), as shown in the protocol in **Figure 2a** and **b** of the manuscript. **b**, The change in DCF fluorescence over 1 minute at baseline (0), over the first (0-60s), second (60-120s), third (120-180s) and fourth minute (180-240s) are shown in *Mybpc3*-KI myocytes in the presence of either vehicle (DMSO; n=11) or CGP (10 µM; n=13; same color-coding as in **a**), respectively. **c-e**, Cardiac myocytes of C57BL/6N mice were stimulated at 1 Hz and then the $Ca^{2+}$ sensitizer EMD 57033 was added to the solution at 0.3 and then 3 µM. Systolic and diastolic sarcomere length (**c**), fractional sarcomere shortening (**d**) and systolic and diastolic $[Ca^{2+}]_c$ (**e**) are indicated.

Error bars indicate the standard errors of the mean (SEM); n-numbers are indicated as the range of numbers of myocytes. $^{+}p<0.05$ unpaired t-test; $^{*}p<0.05$ and $^{**}p<0.01$ (Bonferroni post-test) and $^{\#}p<0.05$ (2-way ANOVA).

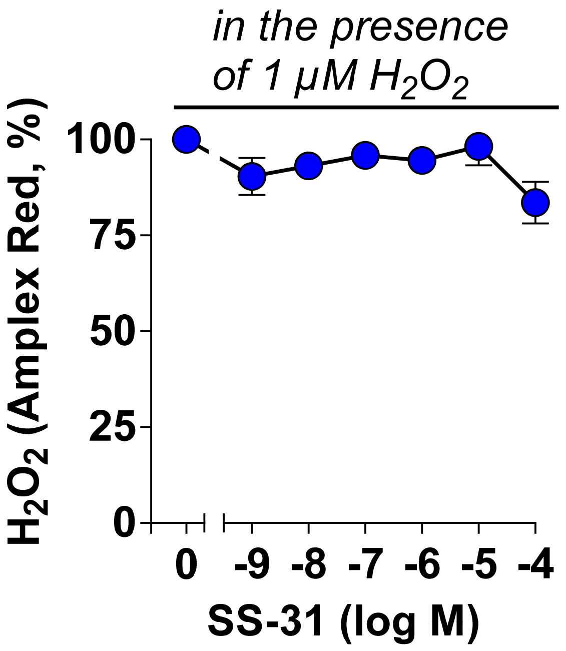


**Supplemental Figure 5:** SS-31 does not directly scavenge $H_2O_2$.

Experiments were performed in a cell free system containing Amplex®UltraRed (Life technologies, Molecular Probes®; 50 µM), horseradish peroxidase (0.5 U/ml), 100 µM $H_2O_2$ and the indicated concentrations of SS-31. Fluorescence of Amplex®UltraRed was recorded in a Tecan GENios Pro Reader at 37°C.

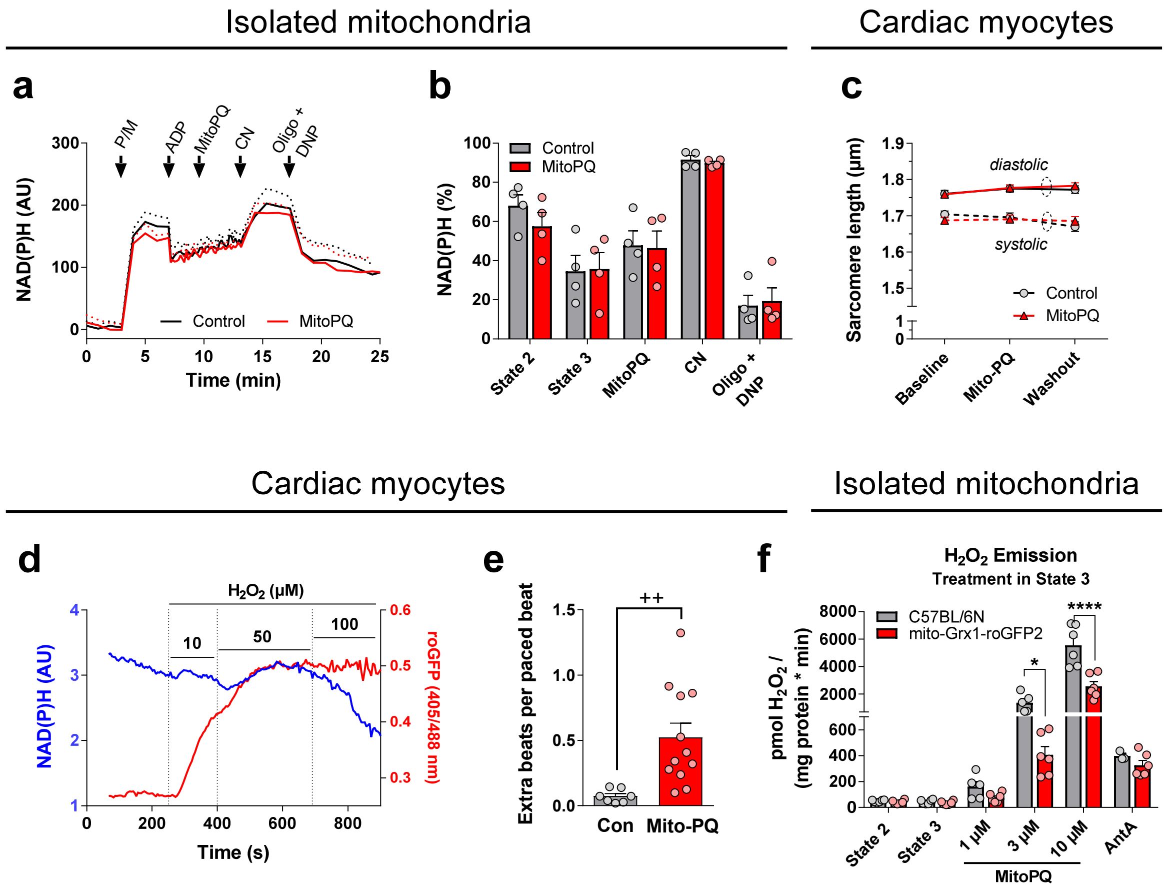


**Supplemental Figure 6:** Effect of Mito-PQ in isolated mitochondria and isolated, intact cardiac myocytes.

Isolated cardiac mitochondria from C57BL//6N mice (n=4) were supplied with pyruvate and malate (5/5 mM) and then exposed to 100 µM ADP to induce state 3 respiration. Mito-PQ (10 µM) was added at the indicated time point. To calibrate the NAD(P)H signal, cyanide (CN; 2.5 µM) for full reduction and subsequently, oligomycin (1.2 µM) and DNP (250 µM) to oxidize NAD(P)H were added. **a**, Original averaged traces and **b**, quantification of NAD(P)H fluorescence after calibration at the indicated conditions. **c,** Isolated cardiac myocytes were exposed to 10 µM Mito-PQ (n=40 cell of n=8 mice) or vehicle (n=39/8) and diastolic and systolic sarcomere length were determined. **d**, Representative example of a cardiac myocyte of a mouse expressing mito-roGFP2-Orp1 exposed to 10-100 µM extracellular $H_2O_2$. The fluorescence of mito-roGFP2-Orp1 (red) was recorded together with NAD(P)H autofluorescence (blue). **e**, Extra beats per regular beats were determined in isolated cardiac myocytes from mice with mito-roGFP2-Orp1 expression in response to vehicle (n=7) or Mito-PQ (n=12). **f**, Isolated cardiac mitochondria from BL/6N mice or mice with mito-Grx1-roGFP2 expression were supplied with pyruvate/malate (5/5 mM), exposed to ADP (100 µM; State 3) and then exposed to 1-10 µM Mito-PQ and finally, antimycin A (AntA; 1.5 µM), and $H_2O_2$ emission determined with Amplex Red (n=3 runs of n=2 mice, respectively, totaling to n=6 cells per condition).

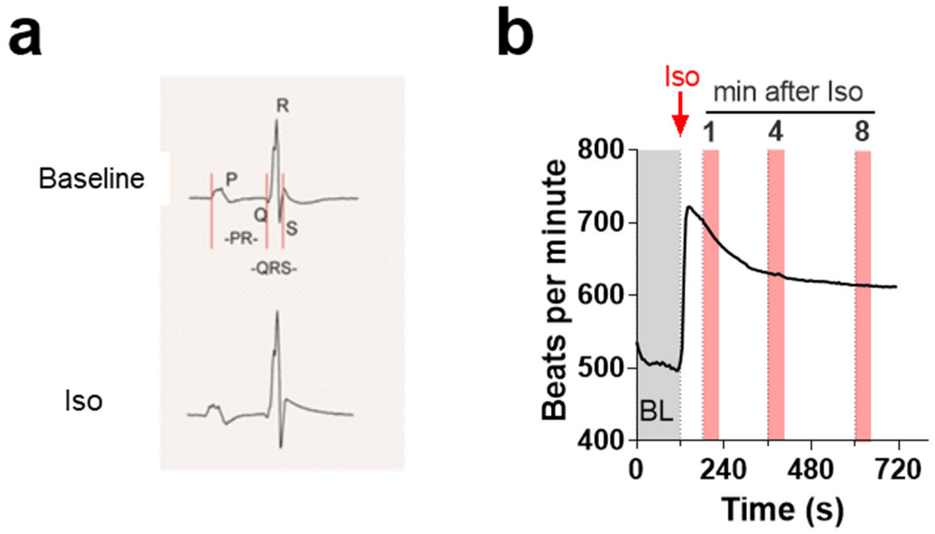


**Supplemental Figure 7:** Protocol to test the effect of SS-31 or vehicle on *Tnnt2*-I79N-induced ECG changes *in vivo*

**a**, Representative QRS complex highlighting how P, Q, R and S were determined and accordingly, PR- and QRS-intervals. **b**, Representative example of heart rate response (beats per minute) after i.p. injection of isoproterenol (Iso, 3 mg/kg). Measurements were performed at baseline before and 1, 4 and 8 min after Iso injection

**Supplemental Table 1**: Arrhythmia scoring system

| Score | Type of Arrhythmia |
|---|---|
| 0 | No PVC, couplet, or VT |
| 1 | < 5 PVC |
| 2 | >/= 5 PVC |
| 3 | couplet |
| 4 | 1 episode of VT |
| 5 | >1 episode of VT |

**Supplemental Table 2:** Electrocardiographic parameters of *in vivo* experiments using Mito-paraquat (MPG).

| **Treatment** | **Vehicle** | **Vehicle** | **0.1mg/kg MPQ** | **0.1mg/kg MPQ** | **0.5mg/kg MPQ** |
|---|---|---|---|---|---|
| 1.5 mg/mg ISO | (-) | (+) | (-) | (+) | (-) |
| Number (n) | 7 | 7 | 8 | 8 | 10 |
| Max Heart Rate (bpm) | 274 ± 31 | 609 ± 30 | 418 ± 24 | 679 ± 19 | 498 ± 36 |
| Max PR (ms) | 40.1 ± 0.17 | 45.6 ± 3.66 | 36.2 ± 1.76 | 62.4 ± 9.13 | 96.3 ± 0.65 |
| Max QRS (ms) | 10.9 ± 0.48 | 13.0 ± 0.77 | 10.3 ± 0.35 | 19.0 ± 1.92 | 17.0 ± 0.65 |
| 3°AV Block | 0/7 | 0/7 | 0/8 | 7/8 | 10/10 |
| Death | 0/7 | 0/7 | 0/8 | 7/8 | 10/10 |
| Arrhythmia Score (Sum) | 0 | 0.3 ± 0.2 | 0 | 8.4 ± 1.9 | 2.6 ± 1.46 |
